# The compact genome of *Giardia muris* reveals important steps in the evolution of intestinal protozoan parasites

**DOI:** 10.1101/870949

**Authors:** Feifei Xu, Alejandro Jiménez-González, Elin Einarsson, Ásgeir Ástvaldsson, Dimitra Peirasmaki, Lars Eckmann, Jan O. Andersson, Staffan G. Svärd, Jon Jerlström-Hultqvist

**Affiliations:** Department of Cell and Molecular Biology, BMC, Box 596, Uppsala University, SE-751 24 Uppsala, Sweden; Department of Medicine, University of California, San Diego, La Jolla, California, USA

**Keywords:** parasite, diplomonad, *Giardia*, streamlined, intestinal colonization, evolutionary biology, horizontal gene transfer, antigenic variation

## Abstract

Diplomonad parasites of the genus *Giardia* have adapted to colonizing different hosts, most notably the intestinal tract of mammals. The human-pathogenic *Giardia* species, *Giardia intestinalis*, has been extensively studied at the genome and gene expression level, but no such information is available for other *Giardia* species. Comparative data would be particularly valuable for *Giardia muris*, which colonizes mice and is commonly used as a prototypic *in vivo* model for investigating host responses to intestinal parasitic infection. Here we report the draft-genome of *G. muris*. We discovered a highly streamlined genome, amongst the most densely encoded ever described for a nuclear eukaryotic genome. *G. muris* and *G. intestinalis* share many known or predicted virulence factors, including cysteine proteases and a large repertoire of cysteine-rich surface proteins involved in antigenic variation. Different to *G. intestinalis, G. muris* maintains tandem arrays of pseudogenized surface antigens at the telomeres, whereas intact surface antigens are present centrally in the chromosomes. The two classes of surface antigens engage in genetic exchange. Reconstruction of metabolic pathways from the *G. muris* genome suggest significant metabolic differences to *G. intestinalis*. Additionally, *G. muris* encodes proteins that might be used to modulate the prokaryotic microbiota. The responsible genes have been introduced in the *Giardia* genus via lateral gene transfer from prokaryotic sources. Our findings point to important evolutionary steps in the *Giardia* genus as it adapted to different hosts and it provides a powerful foundation for mechanistic exploration of host-pathogen interaction in the *G. muris* – mouse pathosystem.

**Author summary:** The *Giardia* genus comprises eukaryotic single-celled parasites that infect many animals. The *Giardia intestinalis* species complex, which can colonize and cause diarrheal disease in humans and different animal hosts has been extensively explored at the genomic and cell biologic levels. Other *Giardia* species, such as the mouse parasite *Giardia muris*, have remained uncharacterized at the genomic level, hampering our understanding of *in vivo* host-pathogen interactions and the impact of host dependence on the evolution of the *Giardia* genus. We discovered that the *G. muris* genome encodes many of the same virulence factors as *G. intestinalis*. The *G. muris* genome has undergone genome contraction, potentially in response to a more defined infective niche in the murine host. We describe differences in metabolic and microbiome modulatory gene repertoire, mediated mainly by lateral gene transfer, that could be important for understanding infective success across the *Giardia* genus. Our findings provide new insights for the use of *G. muris* as a powerful model for exploring host-pathogen interactions in giardiasis.

## Background

Many eukaryotes have evolved from free-living to parasitic lifestyles over evolutionary time, yet parasitism has developed independently in different taxonomic groups and is therefore characterized by many unique features(1,2). Comparative genomics provides an opportunity to investigate the factors of parasitism such as loss of morphological, metabolic and genomic complexity, and consequently reduced evolutionary potential for a free-living lifestyle(2). It can also identify the drivers and consequences of a parasitic lifestyle and generate new testable ecological and evolutionary hypotheses(3).

*Giardia* is a protozoan parasite that non-invasively colonizes the intestinal tract of many vertebrates. The human pathogen, *Giardia intestinalis*, is estimated to cause 300 million cases of giardiasis in the world each year, being a major cause of diarrheal disease(4). Giardiasis is also a problem in domestic animals, and the zoonotic potential of *Giardia* has been highlighted in recent years(5). *In vitro* models of the interaction of *G. intestinalis* with human cells have helped to unravel clues to how *Giardia* causes disease(6–8), such as the importance of, the adhesive disc for attachment(9), flagella for motility(10,11), secreted cysteine proteases for interference with host defenses(12–16), interactions with the intestinal microbiota(6,17), differentiation into cysts for transmission(4,18) and interference with nitric oxide (NO) production(19,20). Despite this progress it remains uncertain whether these *in vitro* models are representative of the natural infection, particularly because animal models of *G. intestinalis* infection have significant limitations. For example, infection of mice, the most commonly used laboratory animals, with human *G. intestinalis* isolates is unreliable and requires manipulations such as antibiotic conditioning(6). *Giardia muris*, one of six recognized species of *Giardia*(21), has been used as a mouse model since the 1960s for exploring the pathogenesis and immunological responses of the mammalian host to infection(22). The availability of knock-out mice and other host-related resources makes *G. muris* a powerful model to investigate host-pathogen interactions(6). The life cycle and infective process of *G. muris* is closely related to infection by *G. intestinalis*(23). Major findings in *Giardia* biology such as flagellar and disc function, cellular differentiation(22,24,25), and immunity(23,26–29) have been pioneered with *G. muris*, and later been shown to be transferable to human *G. intestinalis* infections(29,30). Unfortunately, research on *G. muris* has been hampered by the lack of genome information and gene expression data(5).

Here we describe the draft genome of *G. muris*, representing the first genome of any *Giardia* outside of the *G. intestinalis* species complex. We performed comparative genomics with free-living (*Kipferlia bialata*(31)) and parasitic (*G. intestinalis*(32–34) and *S. salmonicida*(35)) relatives to *G. muris* to determine how *G. muris* may have evolved into an intestinal pathogen of rodents.

## Results

### Genome assembly

We extracted DNA from freshly excysted *G. muris* cysts purified from the feces of infected mice (Fig. S1A) and assembled a high-quality draft genome using sequences obtained by PacBio and Illumina technologies. In addition, we generated RNA-Seq data for gene prediction and gene expression analyses with total RNA extracted from cysts, recently excysted cells (excyzoites), and trophozoites isolated from the small intestine of infected mice (Fig. S1A).

The *G. muris* draft genome consists of 59 contigs spanning 9.8 Mbp, which is notably smaller than the *G. intestinalis* WB genome (12.6 Mbp, Table 1). Most of the genome (9.0 Mbp, 92%) is found on five contigs (>1 Mbp). Of the remaining short contigs (<30 kbp), ten are terminated in telomeric repeats (TAGGG), suggesting they are the terminal points of five chromosomes. The karyotype of *G. muris* was previously shown to consist of four separable chromosomes(36). We hypothesize that our five major contigs represent a total of five chromosomes in *G. muris*, two of which are so close in size (1.290 and 1.297 Mbp) that they were not readily resolved using pulsed-field gels, and are named accordingly from 1 to 5 from largest to smallest in size (Fig. 1). 42 of the 44 small contigs contain ribosomal DNA (rDNA) clusters that encode 28S, 18S, and 5.8S rRNAs. In fact, rDNA clusters make up 2.0% of the total genome, and account for 91.6% of the identified repeats (Methods S1). Half of the contigs terminated by telomeric repeats have adjacent rDNA clusters (Fig. S1B), suggesting that multiple of the *G. muris* chromosomes(36), like those in *G. intestinalis*(37), have long repeats of rDNAs close to the telomeres. In contrast to *G. intestinalis* chromosomes(37), no retrotransposon sequences were found in the telomeric regions and overall very few retrotransposon sequences were detected in the *G. muris* genome.

**Table 1.** Comparison of genome content between *G. muris, G. intestinalis, S. salmonicida and K. bialata*

| Species | *G. muris* | *G. intestinalis* | *S. salmonicida* | *K. bialata* |
| --- | --- | --- | --- | --- |
| **Genome size (Mbp)** | 9.8 | 12.6 | 12.9 | 51.0 |
| **Chromosomes** | 5 (59) | 5 (35) | 9 (233) | ND |
| **(scaffolds)** |  |  |  | (11,564) |
| **G+C %** | 54.7 | 46.3 | 33.4 | 49.4 |
| **Number of protein encoding genes** | 4653 | 4963 | 8067 | 17,389 |
| **Mean / Median protein size (aa)** | 578 / 428 | 635 / 457 | 373 | 333 |
| **\*Mean / Median intergenic size (bp)** | 264 / 37 | 470 / 81 | 421 | 597 |
| **§Coding density %** | 84.5 / 88.6 | 81.5 / 84.7 | 72.1 | ND |
| **Number of introns** | 3 *cis*,5 *trans* | 8 *cis*,5 *trans* | 3 *cis* | 124,912 |
| **tRNA genes** | 68 | 65 | 145 | ND |
| **ASH %** | 0.016 | 0.028 | 0.15 | ND |
| **References** | This study | Ref(32) | Ref(38) | Ref(31) |

\* Mean/Median intergenic distance is based on all RNAs (mRNAs, tRNAs, rRNAs), but not pseudogenized genes.
§ Coding density: First value is based on all RNAs (mRNAs, tRNAs, rRNAs), but not pseudogenized genes; Second value is based on all RNAs including $\psi$ VSP.

**Figure 1.**
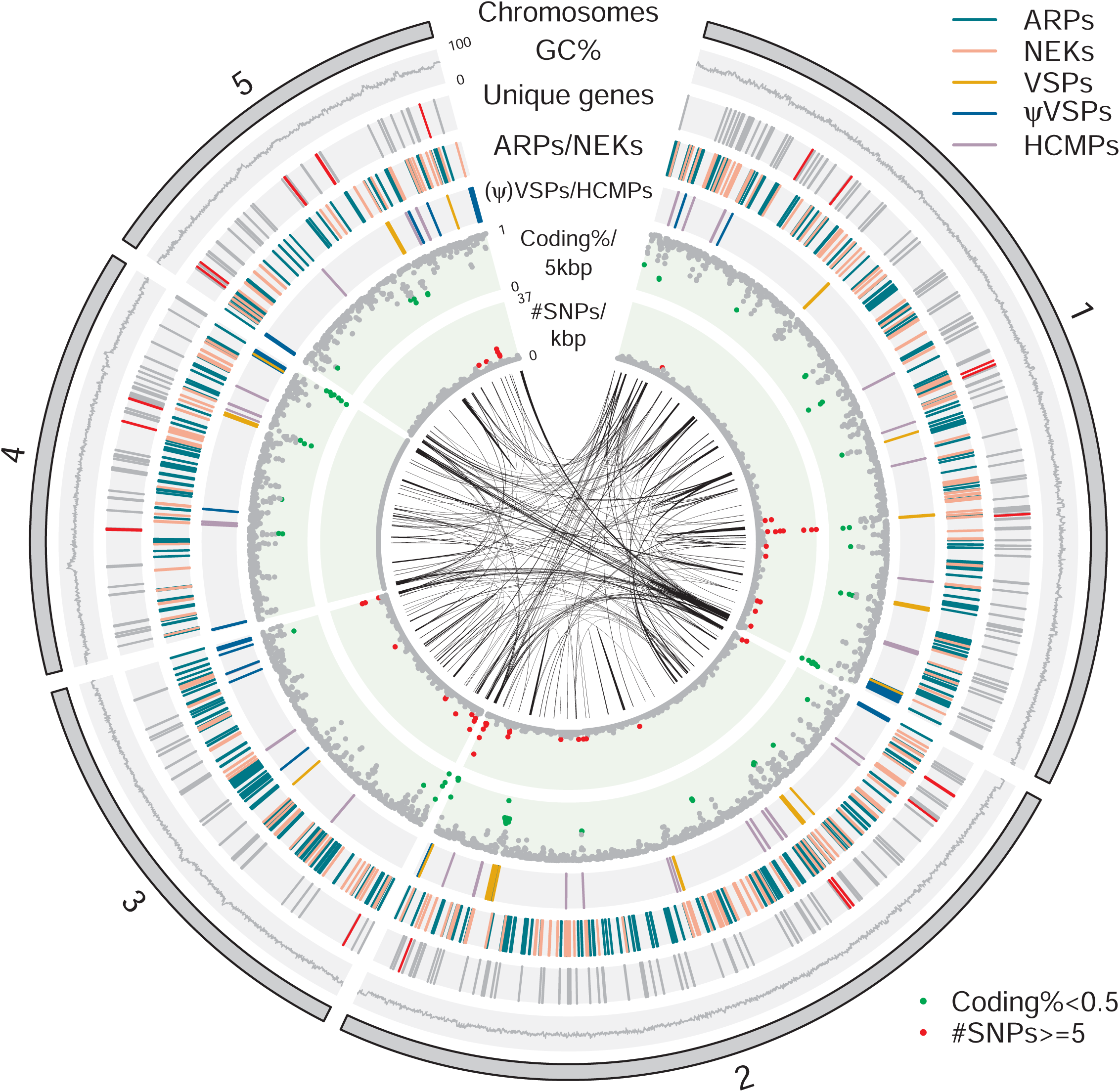
Circular representation of the *G. muris* chromosomes. From outside inward: five chromosomes, GC percent, unique genes (grey) including unique metabolic genes in Table 2 (red), ARPs (greenblue) / NEKs (pink), VSPs (orange) / ΨVSPs (blue) / HCMPs (purple), Coding percent / 5 kbp (green if <= 0.5), # SNPs / kbp >= 5 (red), BLASTN matches with >95% identity and > 1000 bp in size. Circular plot was drawn with circlize(93).

Allelic sequence heterozygosity (ASH) in the assembly was estimated to be 0.016% (Table 1), equivalent to the low level found in the *G. intestinalis* WB genome (0.026%, Table 1). Distribution of ASH along chromosomes showed only weak clustering in certain areas, particularly at the ends of chromosomes (Fig. 1).

**Table 2.**
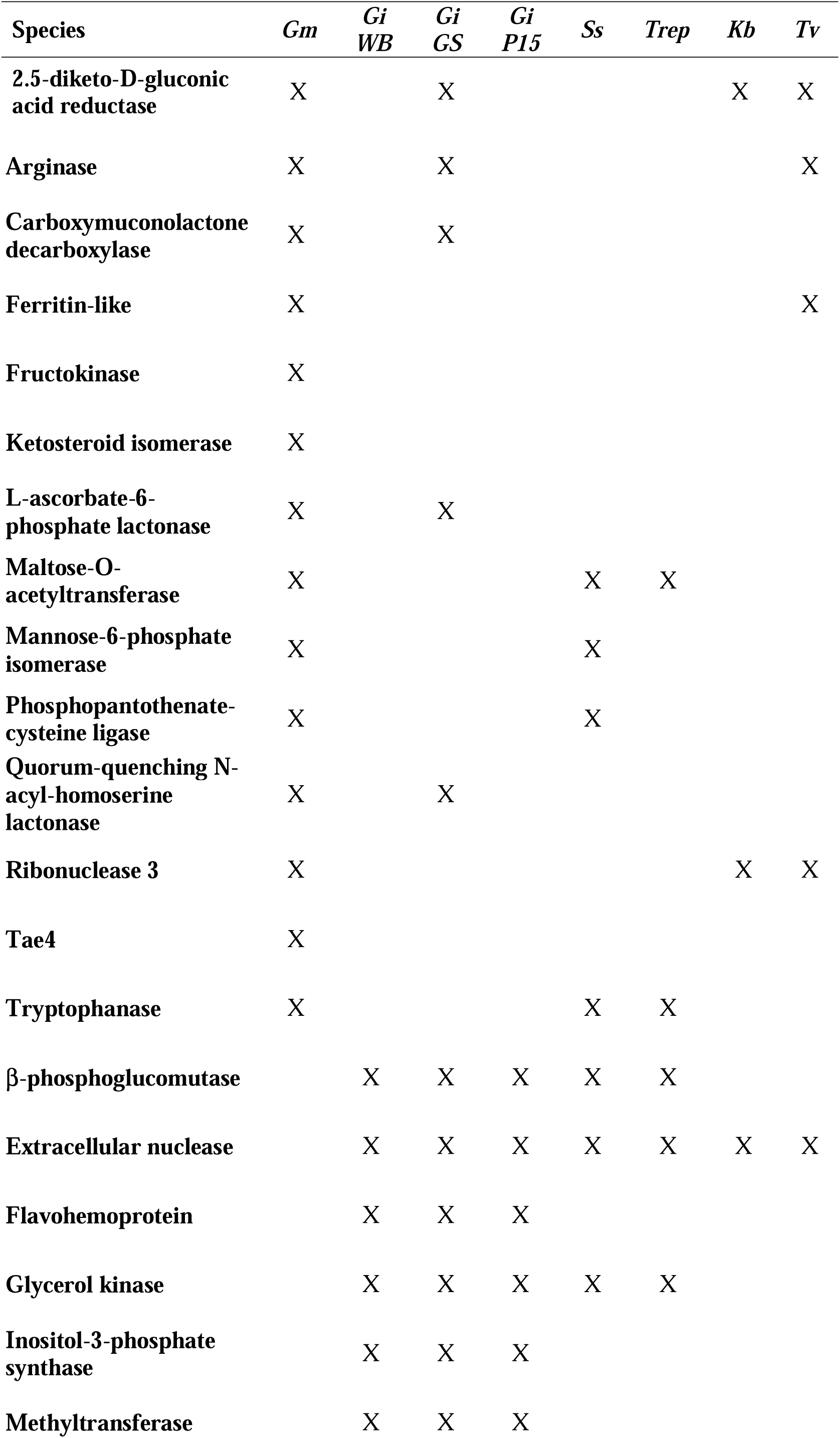
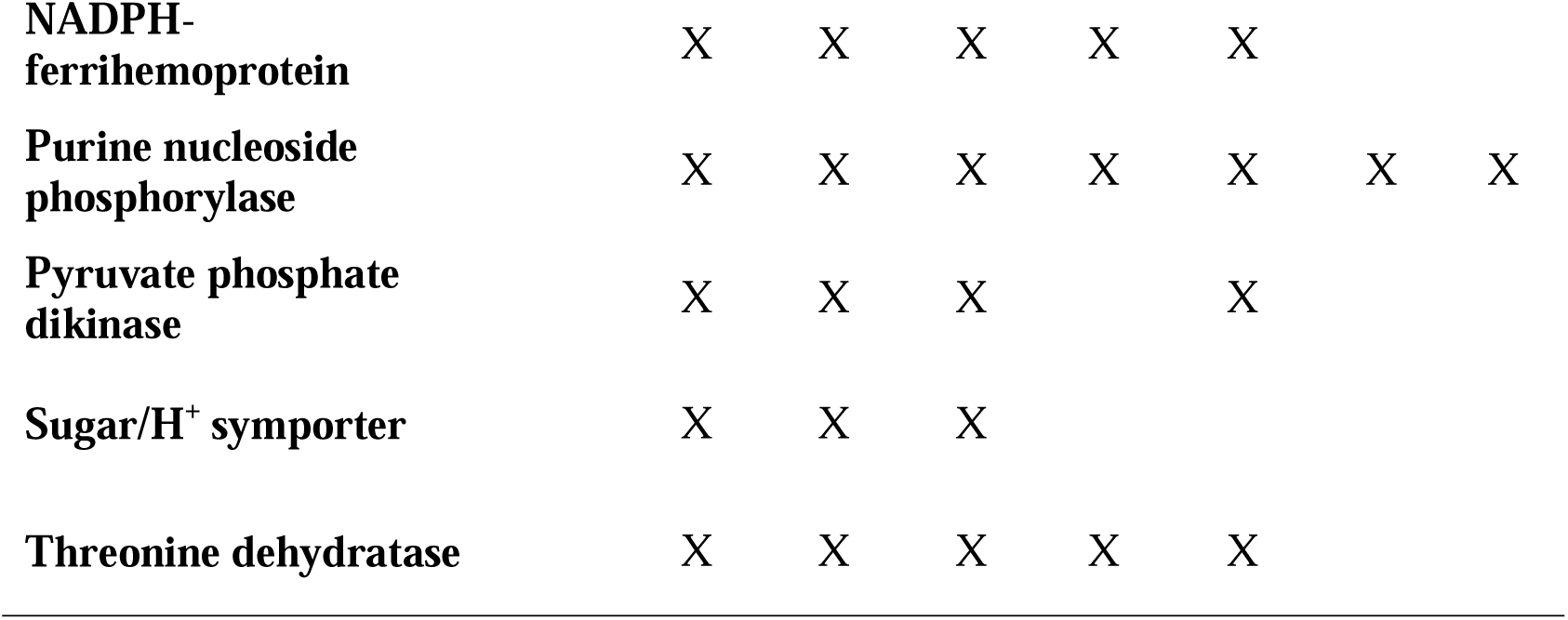
Lateral gene transfers in *G. muris (Gm), G. intestinalis (Gi* WB, *Gi* GS, *Gi* P15*), S. salmonicida (Ss), Trepomonas spp. (Trep), K. bialata (Kb)* and *T. vaginalis (Tv)*.

### Genome streamlining and synteny

Gene prediction and manually curated annotation identified 4653 protein coding genes in *G. muris* (Table 1). This makes 84.5% of the genome coding, counting also the tRNAs and rRNAs (Table 1). Thus, the *G. muris* genome is an example of a very compact eukaryotic genome. Consistent with that, the average intergenic size is 264 bp (Table 1), with a prominent skew towards shorter intergenic regions for a high proportion of genes (median size at 37 bp, Fig. 2C). The compactness of the genome is also illustrated by multiple instances of overlapping genes, with 441 genes (9.5% of all genes) showing an average overlapping size of 21 bp (spanning 1-327 bp) with neighboring genes.

**Figure 2.**
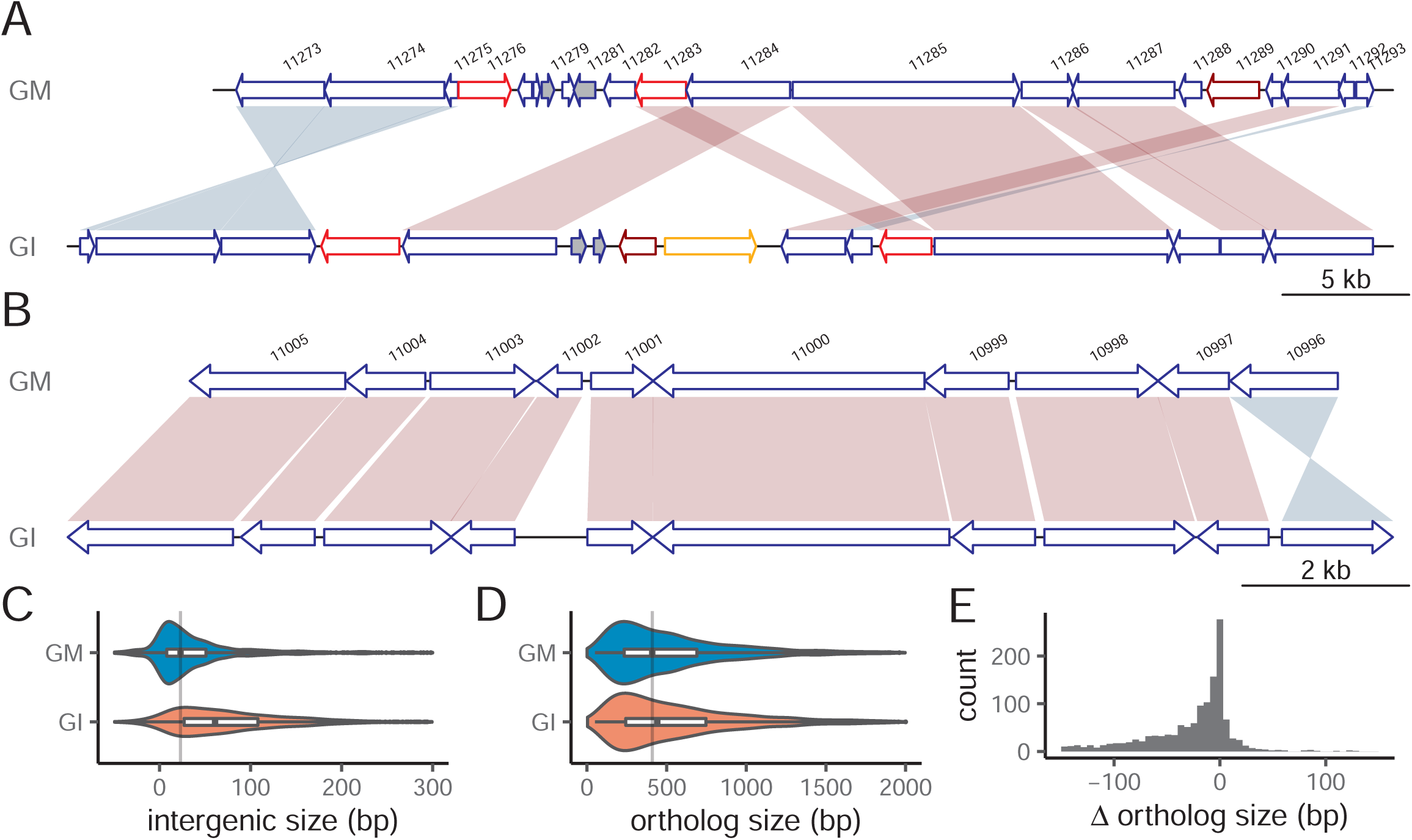
Examples of synteny between *G. muris* and *G. intestinalis*. **A**) A 50 kbp region on chromosome 3 which share synteny to a 58 kbp region on chromosome 5 in WB. Synteny plot was plotted using genoplotR(94). Shades of red and blue represent forward and inverted matches between orthologs. Genes are drawn as arrows in blue. ARPs in red, NEKs in dark red, and VSPs in orange. Dark grey filled genes are unique genes to that genome in comparison to the other. **B**) A 14 kbp region on chromosome 1 which shares synteny to a 16 kbp region on chromosome 5 in *G. intestinalis*. It uses the same color scheme as in A. **C)** Violin plots of intergenic sizes of neighboring positional orthologs of *G. muris* and *G. intestinalis*, and the grey vertical line represents the median intergenic size of *G. muris*. **D**) Violin plots of positional orthologs sizes of *G. muris* and *G. intestinalis*, and the grey vertical line represents the median ortholog size of *G. muris*. **E**) Histogram of the positional ortholog size difference between *G. muris* and *G. intestinalis*.

The *G. muris* and the new improved *G. intestinalis* WB genome (AACB03000000, F. Xu and S.G. Svärd) do not maintain clear chromosomal synteny even though both are assembled as five near-complete chromosomes (Fig. S2). However, local synteny (Fig. 2AB) was obtained among 3,043 one-to-one orthologs (Table S1A) (an average amino acid similarity of 44.7%) shared by the two genomes. Comparing local synteny, it becomes obvious that *G. muris* keeps shorter orthologous gene and intergenic region sizes (Fig. 2BCD).

### Gene regulation

We could not identify any universal, conserved promoter motifs shared by all *G. muris* genes except for an enrichment of A residues around the start codon (Fig. S3A), which resembles observations in *G. intestinalis*(39). The streamlining of the *G. muris* genome was also apparent at the 3’ end of genes where the putative polyadenylation signal, which is similar to the one described in *G. intestinalis*(39), is overlapping with the stop codon for most genes (Fig. S3B). Genes up-regulated early during encystation in *G. intestinalis* have specific promoter elements(40,41). Most of these genes were also identified in *G. muris*, with one notable exception of cyst-wall protein 3 (Fig. 3C). Encystation-related genes in *G. muris* share promoter motifs (Fig. 3A), similar to the Myb binding sites found in *G. intestinalis*(41), suggesting a similar type of regulation. We also noted that the encystation-related genes are among the most highly expressed genes in *G. muris* trophozoites *in vivo* (Fig. S3C, Table S1B), similar to *G. intestinalis* infection in mice(42,43).

**Figure 3.**
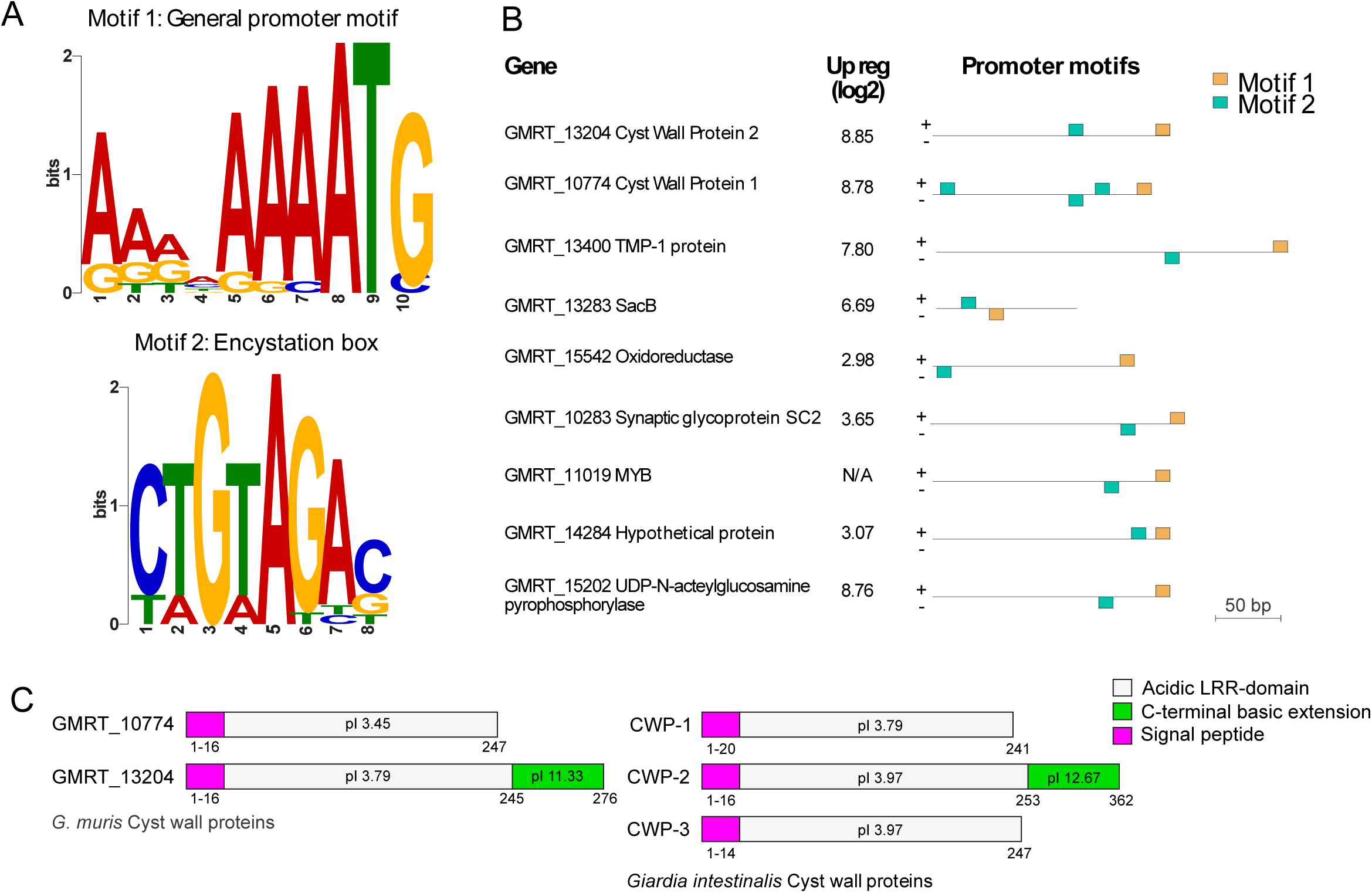
Gene regulation and organization of VSPs in *G. muris*. **A**) Promoter motifs shared by encystation-related genes. Motif 1 (gold in B) represents the general promotor motif positioned directly adjacent to the start codon. Motif 2 (teal in B) resembles the encystation-regulated promoter previously identified in *G. intestinalis* (52). **B)** The distribution and position of motif 1 (gold) and motif 2 (teal) in chosen genes regulated during encystation. **C**) *Giardia* cyst wall proteins. Cyst wall protein 3 is missing in *G. muris*. Signal peptide (pink). Acidic LRR-domain (grey). C-terminal basic extension (green).

Very few genes in *G. intestinalis* contain introns, with only eight known *cis*-spliced and four *trans*-spliced genes (five *trans*-introns)(44–46). Similarly, only three *cis*-and no *trans*-introns were identified in the parasitic diplomonad *S. salmonicida*(35), whereas the free-living fornicate *K. bialata* has on average seven *cis*-introns per protein encoding gene(31). *G. muris* maintains homologs to the eight *cis*-spliced *G. intestinalis* genes, but has only three retained introns (Fig. S4A). All four *trans*-spliced genes in *G. intestinalis* have homologs in *G. muris* with conserved splicing motifs (Fig. S4). Mining genes with similar motifs did not reveal additional intron-containing genes in *G. muris*. Similar to *G. intestinalis*(44) all the *trans*-spliced genes in *G. muris* preserve a similar cleavage motif TCCTTTACTCAA (Fig. S4C) as the RNA processing sequence motif(44). Thus, we observe a reduction of introns in *G. muris*, and *cis*-introns seem to be easier to lose than *trans*-introns.

### VSPs and antigenic variation in *G. muris*

Variant specific-surface proteins (VSPs) in *G. intestinalis* are characterized as cysteine-rich proteins with frequent CXXC motifs and a conserved C-terminal transmembrane (TM) domain followed by a cytoplasmic pentapeptide (CRGKA, Fig. S5A). We identified 265 VSP homologs in *G. muris*. Their C-terminal pentapeptide (GCRGK, Fig. S5A, Table S1C) differed slightly from that in *G. intestinalis*. However, the cysteine and arginine residues in the pentapeptide, which are known to be post translationally modified in *G. intestinalis*, are conserved(47,48). In addition, the preceding 24 aa of the *G. muris* VSP TM domain show conservation to the TM domain of *G. intestinalis* VSPs (Fig. S5A). Most *G. muris* VSPs contain the conserved GGCY motif present in most *G. intestinalis* VSPs (Table S1C). Since *bona fide* VSPs need signal peptides (SPs) at the N-terminus to guide VSPs to the parasite surface, we divided this group into two subgroups; proteins with predicted SPs are called VSPs, whereas VSP proteins without SPs are referred to as pseudogenized VSPs (ΨVSPs). The 26 complete VSP genes (16 unique at 98% identity to each other) are mostly located chromosome-centrally (Fig. 1, Table 3). Seven pairs of VSP genes were identified with identical sequences arranged either as head-to-head (2 pairs) or tail-to-tail (5 pairs) (Table 4). Sequences in between the different VSP pairs resemble NimA (never in mitosis gene a)-related kinase (NEKs), ankyrin repeat proteins (ARPs) and zinc-finger domains. There are also 4 copies of identical VSPs clustering close to the 3’ end of chromosome 2 (Fig. 2) with tandem repeats and sequences resembling NEKs and zinc-finger domains interspersed (Table 4).

**Table 3.** Summaries of gene families within *Giardia*

|  | *G. intestinalis* * | *G. muris* * |
| --- | --- | --- |
| **NEK** | 184 (26) | 216 (23) |
| **ARP-1** | 269 (5) | 298 (6) |
| **ARP-2** | 33 | 86 (16) |
| **ARP-3** | 8 | 33 |
| **VSP** | 133 | 26 |
| **ψVSP** | 208 | 239 |

\* Values in () indicate the number of pseudogenized copies.

**Table 4.** Arrangement of VSP genes in the *G. muris* genome.

| Chr | Geneid<br>1 | Geneid<br>2 | Arrangement | Gene<br>size (aa) | Distan<br>ce (bp) | Genes in between |
| --- | --- | --- | --- | --- | --- | --- |
| 1 | 20512 | 13275 | -> <- | 596 | 2025 | ψARP-2 |
| 1 | 21145 | 21149 | -> <- | 594 | 1658 | ψARP-2 |
| 1 | 13124 | 21374 | -> <- | 620 | 8400 | ψARP-2, alpha-tubulin |
| 2 | 16008 | 21957 | <- -> | 624 | 8794 | NEK, ARP-2 |
| 2 | 22301 | 22304 | -> <- | 523 | 1656 | ψARP-2 |
| 2 | 12920 | 22758<br>(22764,<br>22769) | <- -> | 515 | 10612<br>(6824,<br>8428) | NEK, TRP, Zinc |
| 4 | 24220 | 24228 | -> <- | 623 | 8303 | ψARP-2, alpha-tubulin |
| 5 | 24787 | 24792 | <- -> | 619 | 7158 | NEK, Zinc, ARP-2 |

In contrast to complete VSPs, most ΨVSPs (183 / 239) are found in linear arrays (n=17) in *G. muris*, herein defined as having >3 ΨVSPs genes (Fig. S5BC). Strikingly, nine out of the ten ends of the main contigs have a ΨVSP array at or close to the ends of the chromosomes (telomere-adjacent) containing a total of 131 genes (Table S1C). The only main contig that ends without an array has a cluster of two ΨVSPs close to the chromosome terminus. We found a single ΨVSP array consisting of 12 genes in a chromosome position that was non-telomere adjacent (on chromosome 5) (Fig. 1). The ΨVSP arrays vary in copy numbers (5-23 genes) and the terminal part of the ΨVSP array is always arranged with the tail-end towards the chromosome terminus. The tandem arrangement of the gene arrays suggested that they were generated by gene duplication. Two of the terminal arrays are scrambled and have a shift in the ΨVSP array directionality at the site of an intact VSP (Fig. S5B).

We constructed a phylogeny of all the VSPs and ΨVSPs in *G. muris* to investigate their evolutionary dynamics. The phylogeny revealed relaxed clustering of ΨVSP genes originating in each linear array (Fig. S5C). However, the internal linear array on chromosome 5 represents a noteworthy exception showcasing a very recent gene duplication event. Interestingly, the great majority of full-length VSPs (23/28) are clustered in the phylogeny, including the VSPs in the scrambled ΨVSP arrays, despite these genes being distributed in physically separate chromosomal locations across all five primary scaffolds (Fig. S5C). The few non-clustered VSP genes in the phylogeny that are not part of pairs or clusters are found directly adjacent to ΨVSPs genes. The relaxed clustering of VSPs and ΨVSPs suggested that these genes might be undergoing periodical recombination or gene conversion.

The VSPs and ΨVSPs showed distinct expression patterns. Essentially all the ΨVSP genes were non-transcribed in the three surveyed life-stages (Fig. S5D). VSPs, on the other hand, showed on average higher expression with one or a few loci displaying dominant expression in the different life-stages (Fig. S5D).

There is also another, less characterized VSP-related cysteine-rich protein family in *G. intestinalis*, High Cysteine Membrane Proteins (HCMPs)(49), with 62 members(32). Many are highly up-regulated during interaction with intestinal epithelial cells(50). The HCMPs have several CXXC and CXC motifs, one VSP-like transmembrane domain but with longer C-terminals than in the VSPs(50). The 34 genes matching these criteria in the *G. muris* genome were named HCMPs after the corresponding gene family in *G. intestinalis*. They are found spread-out on the five chromosomes (Fig. 1).

### Multigene families in *G. muris*

The largest multigene families in *G. intestinalis* outside the VSPs and HCMPs are the NEKs(51) and ankyrin repeat containing proteins (Protein 21.1)(32). There are 230 NEKs in *G. muris* (Fig. 1, Table 4), making up 71% of its kinome, slightly more than what was found in *G. intestinalis*(51). We classified ankyrin repeat containing proteins further into three groups. The ankyrin repeat protein-1 (ARP-1) with only ankyrin repeats, ARP-2 with ankyrin repeats plus zinc finger domains, and ARP-3 with both ankyrin repeats plus domains other than zinc finger domains. The NEKs and the different classes of ARPs are scattered throughout the chromosomes without obvious clustering (Fig. 1). A phylogenetic analysis revealed that 79 of the NEKs are conserved as 1:1 orthologs between *G. muris* and *G. intestinalis* (Fig. S6A). Each species has one massively expanded cluster of NEKs with *G. muris* having the largest with 104 members and the one in *G. intestinalis* having 79 members. ARPs show a similar evolutionary stability with 132 conserved 1:1 orthologs between species (Fig. S6B). *G. muris* shows a major species-specific expansion of 91 genes whereas the largest expanded clusters in *G. intestinalis* amounts to two groups of 15 genes each. The partly shared domain-structure of NEKs and ARPs prompted us to investigate their relationship by a network analysis employing reciprocal blastp (1e-05 cutoff). The two groups are not recovered as clearly separated clusters but form partially overlapping networks, indicating that there might be recombination or gene conversion in between (Fig. S6C). The larger numbers of genes in these multigene families in *G. muris* compared to *G. intestinalis* and their evolutionary dynamics are intriguing given the otherwise streamlined features of the *G. muris* genome, perhaps suggesting that they have unique roles in adaptation to their murine hosts.

### Virulence factors in *Giardia*

*G. intestinalis* is not known to possess classical virulence factors, such as enterotoxins, but several genes are important for colonization of the host and thus for pathogenesis. These include genes for motility(10), the adhesive disc for attachment(9), secreted cysteine proteases that can degrade host defensive factors(12,13), and cysteine-rich surface protein like the VSPs(52) and the HCMPs(49) that undergo antigenic variation. The cytoskeletal protein repertoire in *G. muris* is very similar to *G. intestinalis* apart from several fragmented alpha-tubulins (3 complete genes with homologs in *G. intestinalis* and 9 incomplete gene fragments). The adhesive disc is a unique cytoskeletal structure of *Giardia* parasites essential for attachment of the trophozoite in the small intestine, but is missing in other fornicates(9). The first detailed studies of the adhesive disc were performed on *G. muris* trophozoites(24), but more recent work has mostly focused on *G. intestinalis*. The vast majority (82 of 85) of *G. intestinalis* disc proteins(9) were also identified in *G. muris* (Table S1D); 12 were NEK kinases and 27 were ARP-1 proteins. Two of the three *G. intestinalis* disc proteins not found in *G. muris* (Table S1D) localize to a structure in the *G. intestinalis* disc on top of the ventral groove, but this structure is missing in the *G. muris* disc(22), suggesting functional disc differences. Many of the disc proteins that are immunodominant during *G. intestinalis* infections (e.g. alpha-giardins, beta-giardin, SALP-1, alpha-and beta-tubulin(53) are highly expressed (here defined as >500 FPKM) in *G. muris* trophozoites in the small intestine (Table S1B).

Proteases are important virulence factors in many pathogens and cysteine-protease activities have been suggested to play a role in *Giardia* virulence(4,13). We identified 81 proteins classified as proteases in the *G. muris* genome, compared to 96 proteins identified in an identical search in *G. intestinalis* WB (Table S1E). The largest family of proteases in *G. muris*, with 15 members, are papain-like cysteine proteases (C1A family). This protein family is also the largest protease group in *G. intestinalis* with 21 members(54,55). Several conserved groups of proteases were found to have been present in the ancestor to *Giardia* and *Spironucleus*, although we also found evidence for lineage-specific gene loss and expansion in *G. muris* (Fig. S7). The most highly expressed cysteine protease of *G. muris in vivo* is the closest homolog to the highest expressed protease in *G. intestinalis* WB(55).

### Metabolic pathways in *G. muris*

Our metabolic reconstruction identified 95 metabolic pathways in *G. muris* compared to 98 pathways detailed in *G. intestinalis* WB; four pathways were unique in *G. muris* and eight in *G. intestinalis* WB. Even though the overall metabolism is highly similar between *G. muris* and *G. intestinalis* WB, the genes for several specific enzymes and their putative reactions show distinct differences. Thus, 18 unique reactions (14 enzymes) were predicted in *G. muris* and 25 in *G. intestinalis* WB (10 enzymes) (Table 2). Several of these unique proteins showed moderate-high identity to prokaryotic proteins (Table S1F). Five of these prokaryote-like genes are present in the genome of assemblage B strain *G. intestinalis* GS.

The potential utilization of carbohydrate sources for glycolysis is different in *G. muris* compared to *G. intestinalis*. Fructokinase and mannose 6-phosphate isomerase enable *G. muris* to use fructose and mannose 6-phosphate unlike *G. intestinalis*. In both cases, the enzymes were acquired from bacteria via lateral gene transfer (Table S1F, Figure S8AB). Curiously, the *G. muris* gene for mannose 6-phosphate isomerase is found next to a bacterial transcriptional regulator/sugar kinase gene. Phylogenetic analyses show that *G. muris* mannose 6-phosphate isomerase and the transcriptional regulator/sugar kinase gene clusters deep in the Bacteroidetes group. This gene arrangement is observed in bacteria of the genus *Alistipes*, whose genomes harbor the most similar homologs, supporting the notion that a single event of lateral gene transfer best explains the origin of these genes in *G. muris* (Figure S8BC). Both genes have the A-rich initiator that precedes the start codon in most *G. muris* genes and are expressed in *G. muris* trophozoites (Table S1B).

The utilization of glycerol for ATP synthesis via glycerol kinase has been suggested in *G. intestinalis* upon depletion of primary carbon sources(56). This enzyme is found in both *Spironucleus* and *Trichomonas* but has been lost in *G. muris*. Another notable metabolic difference to *G. intestinalis* WB is the lack of pyrophosphate-dependent pyruvate phosphate dikinase (PPDK) that leaves a single, less energy efficient, route from phosphoenol pyruvate to pyruvate via pyruvate kinase in *G. muris*.

*G. muris* is predicted to synthesize coenzyme A from pantothenate, employing a bifunctional phosphopantothenoyl decarboxylase-phosphopantothenate synthase. The same pathway is described in *S. salmonicida*(35), but the complete pathway is missing in *G. intestinalis* (Table 2).

As in all other studied metamonads, *G. muris* encodes the arginine dihydrolase (ADH) pathway that enables the use of arginine as an energy source and at the same time, reduces the available free arginine in the environment, preventing nitric oxide (NO) production in host cells(57). NO efficiently kills *G. intestinalis* trophozoites and the main scavenger enzyme for NO in *G. intestinalis* is Flavohemoprotein(58), which is lacking in *G. muris* (Table 2). Arginine is an important modulator of virulence in many infectious organisms since it interferes with NO production(59). *G. muris* encodes arginases which converts arginine directly to ornithine and urea. Arginases are present in *G. intestinalis* GS and *T. vaginalis*, representing an ancestral acquisition in Metamonada followed by subsequent losses in *Spironucleus* and *G. intestinalis* assemblage A and E (Table 2 and Fig. S8D).

We noticed that several of the bacterial derived genes that are shared with *G. intestinalis GS* are clustered together on the chromosomes in *G. muris* in highly dynamic genomic regions. For example, arginase genes are found in a four-gene genomic region that is present in two adjacent copies in chromosome 2, two on chromosome 1 and one on chromosome 5 (dark grey filled arrows, Fig. 4A). Intact arginase genes are only found on chromosome 2 adjacent to the genes encoding 2,5-diketo-D-gluconic acid reductase. All the duplication events have occurred between ARPs (red arrows) and NEKs (dark red arrows). The homologous regions on the three chromosomes likely originated via two duplication events.

**Figure 4.**
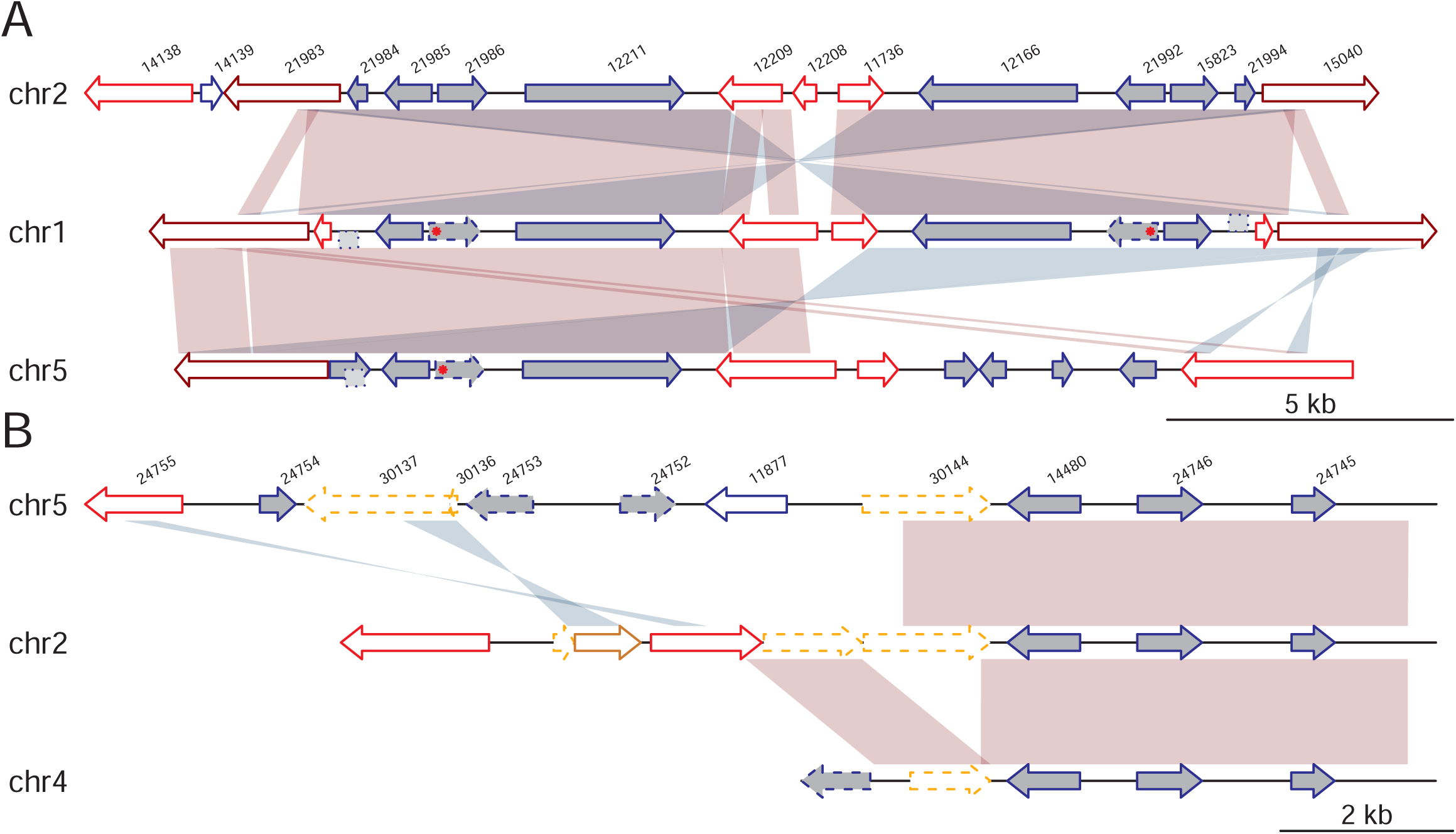
Synteny plot of two duplicated regions in the genome. **A, B)** Shades of red and blue represent forward and inverted matches between neighboring sequences. ARPs are drawn in red, NEKs in dark red and VSPs in orange. Pseudogenized genes are drawn in dashed lines. Dark grey filled genes are unique genes to *G. muris* in comparison with *G. intestinalis*. Point mutation in arginase is marked in red asterisk. Homologous sequences that are not annotated in the genome are drawn in dashed box on sides of the backbone grey line. Genes discussed in the paper: 21985 and 21992 encode the intact arginase genes, 21986 and 21992 encode the enzyme 2,5-diketo-D-gluconic acid reductase, whereas 21984 and 21994 are hypothetical proteins in A; 14480 encodes 2,5-diketo-D-gluconic acid reductase, 24746 encodes CMD and 24745 encodes ketosteroid isomerase-like protein in B.

Nucleotide substitutions have accumulated since the duplication events and in-frame stop codons (marked by red asterisk in Fig. 4A) have rendered three of the arginase genes pseudogenized. The small ORFs sitting at the other side of arginase are hypothetical proteins and have similar homologs in other parts of the genome, but the sequences of their duplicated homologs in those regions are pseudogenized (dotted light grey block in Fig. 4A). Both genes have their closest relatives in bacteria, even though the genes can be found in other eukaryotes (Fig. S13E). Phylogenetic analyses indicate the genes might have been transferred from different bacterial donors multiple times into different eukaryotic lineages (Fig. S13E).

A second, distantly related 2,5-diketo-D-gluconic acid reductase gene copy (Fig. 4B, Fig. S8F) in the genome is present in three different genomic locations together with two other enzymes, carboxymuconolactone decarboxylase (CMD) and ketosteroid isomerase-like protein. All three genes, constitute putative lateral gene transfers (Fig. S8GH). This three-gene region is close to ΨVSPs (orange arrows) and ARPs.

### Interaction with the intestinal microbiota

*Giardia* trophozoites colonize the intestinal lumen where they can potentially interact with other intestinal microbes. Although little is functionally known about such interactions and their consequences for parasite survival, four proteins encoded in the *G. muris* genome could play a role in these interactions.

Bactericidal/permeability-increasing (BPI) proteins are innate immune defense proteins that bind to lipopolysaccharide and display potent killing activity against gram-negative bacteria by increasing membrane permeability. Beyond this basic function BPI proteins might also act as effectors in controlling mutualistic symbioses(60). Homologs of BPI proteins are found in *G. muris* and *G. intestinalis*(61), but it remains to be determined if they have anti-microbial activity.

*G. muris* encodes tryptophanase, an enzyme that metabolizes tryptophan to pyruvate with concomitant release of indole and ammonia. While pyruvate can be utilized in energy metabolism, indole and its metabolites have been shown to affect gut microbiota composition, possibly by interfering with quorum-sensing systems, and might be able to influence host health(62). Phylogenetic analysis of this protein showed that this enzyme represents an ancestral acquisition in diplomonads with subsequent loss in *G. intestinalis* (Fig. S8I).

Two more proteins with potential importance for microbiota interactions are encoded in *G. muris*: Tae4 and quorum-quenching N-acyl-homoserine lactonase. The Tae4 proteins are wide-spread amidases that were first described in association with the T6SS system effector Tae4 in *Salmonella* Typhimurium(63). The Tae4 proteins degrade bacterial peptidoglycan by hydrolyzing the amide bond, γ-D-glutamyl-mDAP (DL-bond) of Gram-negative bacteria(63), and is required for interbacterial antagonism and successful gut colonization by *S*. Typhimurium(64). Quorum-quenching N-acyl-homoserine lactonase degrades N-acyl-homoserine lactone, a molecule used by both Gram-positive and Gram-negative bacteria, for quorum sensing(65). Our phylogenetic analysis supports the lateral acquisition of both genes (Fig. S8JK). While Tae4 has been a recent acquisition in *G. muris*, quorum-quenching N-acyl-homoserine lactonase was present in the common ancestor of *G. muris* and *G. intestinalis* and lost in *G. intestinalis* WB and P15 (Fig. S8K).

## Discussion

Our data shows that *G. muris* has an even more compact genome than *G. intestinalis*, whose genome is already known to be highly streamlined(32). Genome compaction via reduction of mobile or repetitive elements have been seen in other eukaryotic parasites(1,66). *G. muris* appears to fall into this category as it encodes no known classes of mobile elements and repetitive elements are mostly confined to telomeric contexts. The shortness of intergenic regions in *G. muris* ranks among the most extreme recorded for any eukaryote, even shorter than Microsporidia which are known as the most compact and reduced eukaryotic genomes(67). The global synteny map of *G. muris* to *G. intestinalis* indicates many frequent small-scale genome rearrangements that often favors a more efficient gene packing in *G. muris* thus allowing shorter intergenic regions. This evidence of gene shuffling and the fact that there is very little evidence of genome degradation would argue for optimization of growth as the driving force of *G. muris* genome streamlining.

*G. muris* trophozoites have not been grown axenically *in vitro*, which has hampered exploration of its genome, gene regulation and metabolism(5,21), and has limited the use of *G. muris* as an *in vitro* model system for the human parasite *G. intestinalis* and other intestinal protozoan parasites(5). We identified several metabolic differences between *G. muris* and *G. intestinalis* that might indicate avenues to successful strain axenization. Most of these differences are represented by instances of lateral gene transfer of metabolic genes or losses thereof in either *G. intestinalis* or *G. muris. G. intestinalis* isolates are typically poor at infecting mice. Despite this, the assemblage B isolate GS, which shares more metabolic enzymes with *G. muris* than the assemblage A isolate WB, is better able to establish infection in mice than WB. This suggests that the shared metabolic capacity of *G. muris* and *G. intestinalis* GS enables survival in the murine intestinal tract. Additionally, *G. muris* might be able to interact or interfere with intestinal Gram-positive and Gram-negative bacteria and this could be a key to establish successful infections. *G. intestinalis*, which lacks some of the putative microbiome modulators, such as Tryptophanase and Tae4, is dependent on reduction of the small intestinal microbiota in order to efficiently infect mice(68). *G. muris* is cleared from the murine host by secretory IgA(26), whereas the role of IgA in anti-giardial defense is less clear for *G. intestinalis* GS(26,69). We speculate that *G. muris* is more resistant to elimination by innate factors such as competition with the normal microbiota, or host production of reactive oxygen species and/or NO, whereas GS is more sensitive to innate factors and eliminated much faster within 1-3 weeks (while *G. muris* clearance requires 4-8 weeks). Future insights into the importance of innate factors in *G. muris* infection should be facilitated by the availability of the complete genome sequence.

Sub-telomeric regions in parasitic protozoa often contain arrays of expanded gene families that are under positive selection by the immune system(66). The relaxed evolutionary pressure offered by keeping pseudogenized copies of surface antigens might be an advantage for *G. muris* that allows genetic drift and recombination to drive rapid and stealthy diversification, thus avoiding elimination by adaptive immune defenses. It was previously reported that *G. muris* is capable of antigenic variation and encodes VSP genes with high similarity and conserved structural features (CRGKA pentapeptide) to those *G. intestinalis*(70). We failed to identify close homologs to the previously sequenced *G. muris* VSPs in our *G. muris* genome. The linear ΨVSP arrays in *G. muri*s have previously been described in *G. intestinalis*(71). Our phylogenetic analyses of *G. muris* VSPs and ΨVSPs revealed evidence of recombination or segmental gene conversion, as previously demonstrated in *G. intestinalis*(71). However, we recognized two clear differences in the VSP repertoire in *G. muris* and *G. intestinalis*. First, *G. muris* encodes a low number of intact VSP loci that are located internally on the chromosomes. Second, the ΨVSP arrays are almost exclusively telomere adjacent, as opposed to *G. intestinalis* where this tendency is not apparent(71). These aspects of the *G. muris* VSP repertoire resemble the antigenic variation systems of *Pneumocystis* spp. and *Trypanosoma brucei* (72,73). Despite clear mechanistic differences, all these systems have converged on having large reservoirs of mostly telomeric positioned, arrayed genes that are transcriptionally silent and are sources for recombination and gene conversion into expression sites.

The function of ankyrin repeat proteins and NEK kinases remains mostly unknown. They represent, together with VSPs, the most dynamic protein families in the *Giardia* genomes(51). The *Giardia* NEK kinases lack transmembrane domains and have been suggested to target and localize to different intracellular structures with their ankyrin repeats(51) and many of the *G. intestinalis* NEK kinases localize to cytoskeletal structures, including the flagella and adhesive disc(9). Rearrangements and duplications in the *G. muris* genome are frequently associated with these large gene families (Fig. 2A and Fig. 4), indicating they might serve as anchoring-points for recombination.

Lateral gene transfer is an important shaping factor in the evolution of metabolism in protists(74). The origins of laterally transferred genes in *G. muris* are here inferred to be by prokaryotic sources that are members of the gastrointestinal flora, in agreement to previous observations(75). Most of the putative differences in metabolic potential in the *Giardia* genomes are attributable to lateral gene transfers, either by lineage specific gene gain or loss. For example, the ability to utilize mannose has been introduced from bacteria of the genus *Alistipes* via lateral gene transfer. This event is supported by phylogenetic reconstruction, shows a high degree of sequence conservation (>70% at the amino acid level) and displays maintained gene order to the one seen in the closest related bacterial lineages (Figure S8B). Interestingly, several lateral gene transfers were found clustered in amplified areas of the *G. muris* chromosome. Curiously, *G. intestinalis* assemblage B also maintains clustered copies of arginase and 2,5-diketo-D-gluconic acid reductase, while these genes have both been lost in the *G. intestinalis* assemblage A and E lineages.

Anti-microbial peptides of several classes, such as defensins and trefoil-factor 3, are up-regulated in the small intestine of *G. muris* infected mice(29). Secreted cysteine proteases from *G. intestinalis* have been shown to be able to degrade defensins(12). We detected prominent expression of several cysteine proteases in *G. muris*. The protease with the highest expression is suggested to have a role in encystation and excystation(55). Its *G. intestinalis* homolog is up-regulated and secreted during interactions with human intestinal epithelial cells(8) and it cleaves chemokines, tight junction proteins and defensins(12,76). Thus, this is most likely also an important virulence factor in *G. muris*.

Our results from this study are summarized in a model of the evolution of *Giardia’s* virulence traits in Figure 5. A number of characters important for *Giardia*’s ability to infect the intestine of mammals are pre-parasitic inventions (such as modified mitochondria and differentiation into transmissive cysts, Fig. 5) and some are found in all diplomonad parasites (e.g. loss of metabolic functions, streamlined microtubular cytoskeleton, expansion of gene families like ankyrins and cysteine proteases and loss of introns, Fig. 5). *Giardia* specific innovations include the adhesive disc for attachment, VSPs and High Cysteine Membrane Proteins for antigenic variation and mitosomes involved in Fe-S complex synthesis (Fig. 5). Whereas some are only found in *G. muris* (e.g. metabolic genes involved in microbiota interactions, Fig. 5), suggesting adaptation to the intestinal environment of mice. Our data shows that the environment in the host’s intestine, most of all the immune system and the microbiota, apply selective pressure for changes in the genome, metabolic potential and the parasite surface proteome.

**Figure 5.**
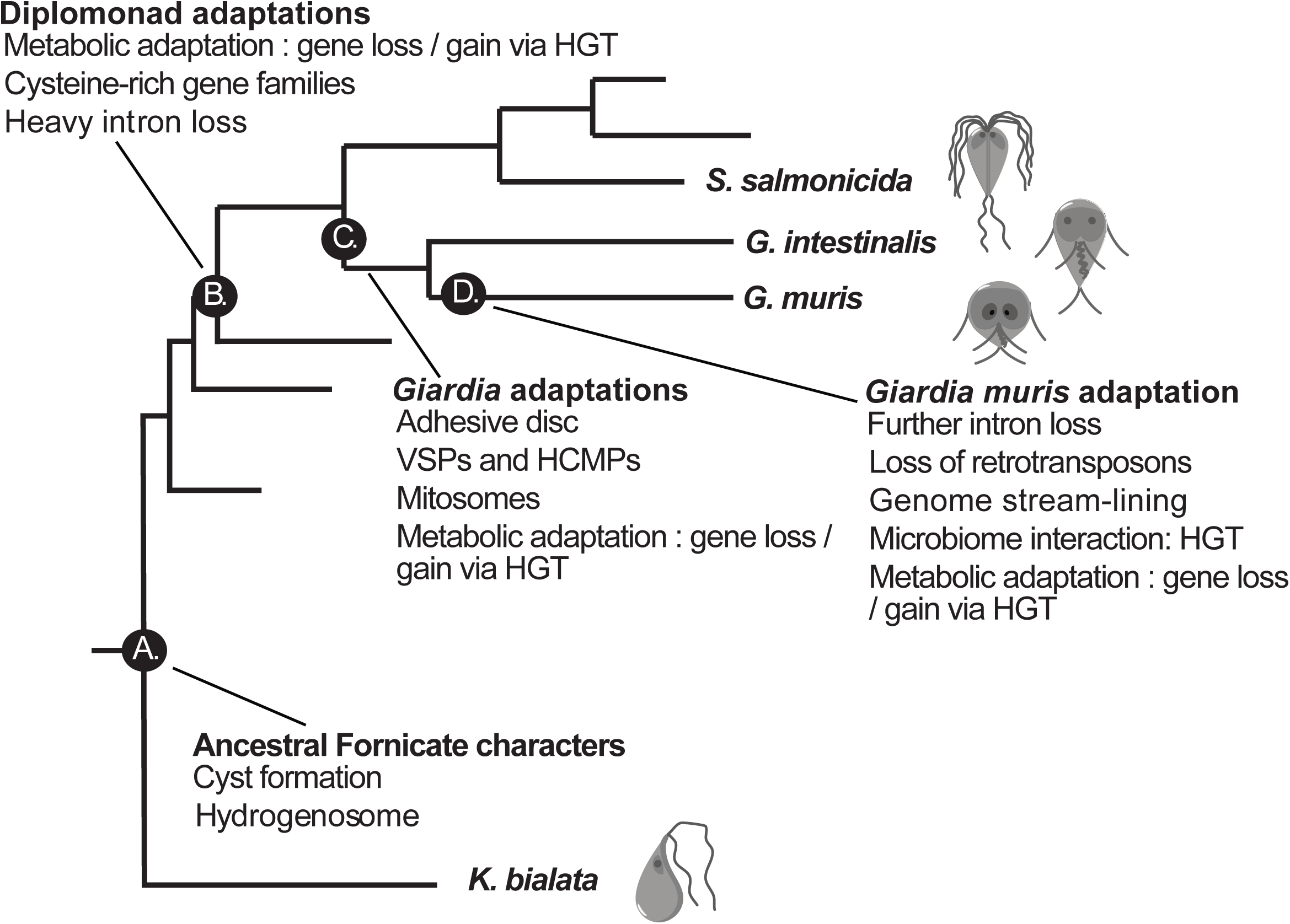
Model of the evolution of virulence traits in *Giardia* parasites. A set of important diplomonad evolutionary innovations and their chronology is depicted at relevant phylogenetic nodes. A) Free-living fornicate ancestor. B) Diplomonad ancestor. C) *Giardia* ancestor. D) *Giardia muris*.

## Materials and methods

### Cell preparation and nucleic acid extraction

4.5×10^7^ muris trophozoites (day 7 post infection) were collected from small intestines of three C57 mice, washed once in PBS and pellet frozen at −80°C (Biosample SAMN11231832). Viable cysts of *G. muris* isolate Roberts-Thomson passaged through mice were obtained from Waterborne Inc. These cysts had been purified from fecal material using Percoll and sucrose gradients (Biosample SAMN11231833). DNA and RNA were extracted from 1×10^7^ cysts using standard methods. 1×10^8^ cysts were excysted according to the procedure in Feely et al.(77) (Biosample SAMN11231834). RNA was purified from cell material equivalent to 1×10^7^ cysts. DNA for long-read sequencing was prepared from the remaining cysts as described in Methods S1.

### Sequencing, assembly and annotation

Total genomic DNA was sequenced using both Illumina MiSeq and PacBio RS II sequencers. The stranded transcriptome mRNA and the miRNA libraries were sequenced with Illumina HiSeq 2000 system. The RNA samples extracted from excysted cells and cysts prior excystation were prepared using the TruSeq stranded mRNA sample preparation kit and sequenced by HiSeq 2500. PacBio long reads were assembled de *novo* using the SMRT Analysis (v2.3.0) pipeline. A detailed description of genome sequencing, assembly, annotation, synteny analyses and RNA-Seq is available in Methods S1.

### Pathway analysis

The metabolic pathways of *G. muris* and *G. intestinalis* WB were predicted with a combination of BlastKOALA(78) implemented in KEGG(79), Pathway Tools v21.5(80) and GiardiaDB(81). The different predictions were combined and manually curated under Pathway Tools(80). Pathway Tools function pathway hole filler(82) was used to further complete the pathway, and transport inference parser(83) was used to infer transport reaction(s) for transporters which were then verified with Conserved Domain databases(84).

### Phylogenetic analysis

*G. muris* sequences were used as queries to retrieve at least 5000 hits with e-value <0.001, using BLASTP against the nr database and the organism-specific proteomes. The datasets were aligned in the forward and reverse orientation using MAFFT v6.603b(85) and PROBCONS v1.12(86). The four resulting alignments were combined with T-COFFEE(87) and trimmed by BMGE v1.12(88). Maximum Likelihood (ML) trees were computed using IQtree v1.6.5(89) under LG4X substitution model(90). Branch supports were assessed using ultrafast bootstrap approximation (UFboot)(91) with 1,000 bootstrap replicates and 1,000 replicates for SH-like approximate likelihood ratio test (SH-aLRT)(92). A detailed description of the phylogenetic analyses is available in Methods S1.

## Supporting information

**Figure S1. A)** *G. muris* cells propagate as trophozoites in the mice small intestine. The cell trophozoites undergo encystation into cysts and are shed in the feces. The cysts are prompted to excyst as they encounter the conditions of the digestive tract. The excyzoites transition into the vegetative trophozoites. Cell material for genomic DNA was harvested from *G. muris* cysts from infected mice. **B)** Sub-telomeric regions in *G. muris*. Sequence motifs in the 10 unitigs terminating in (TAGGG)^n^ telomeric sequences (purple). Ribosomal DNA sequences (black). Satellite-like repeated sequences (green).

**Figure S2. Chromosome-scale synteny between *G. muris* and *G. intestinalis* WB**. *G. muris* chromosomes are color coded. The links between the two genomes representing sequence similarity identified by MUMmer promer(95), are colored after the *G. muris* chromosomal color. Circular plot was drawn with circlize(93).

**Figure S3. Sequence logo around start codon (A) and stop codon (B) in *G. muris*. C) *G. muris* gene expression from *in vivo* trophozoites**. Genes showing top-ranked FPKM ratios (log2 (Trophs/Cysts)) between *in vivo* derived trophozoites and cysts (cut-off value ≥4). The value is indicated in the teal circle. An orange dot indicates that the ortholog of this gene is upregulated (>2 fold) in *G. intestinalis* at 7 hrs, 12 hrs or 22 hrs into encystation (40).

**Figure S4. *Trans*-and *cis*-introns in *G. muris*. A)** Aligned *trans*-and *cis*-splice junctions in *G. muris*. **B)** Splice-site associated motifs in *trans*-spliced exons. Base-pairing stems are underlined. Bases in red denote the predicted cleavage motif. **C)** Consensus alignment of the cleavage motifs in *G. muris*. **D-E)** Predicted secondary structures of **D)** *cis*-spliced and **E)** *trans*-spliced genes. The secondary structure was predicted using the mfold webserver using default settings and manually curated. GU-rich stretches and cleavage motifs are boxed.

**Figure S5. A) Conservation of the VSP C-terminal in *G. muris* and *G. intestinalis***. Sequence logos of conserved motifs in the C-terminal of VSPs in *G. muris* (n=25) (upper panel) and *G. intestinalis* WB (n=183) (lower panel) were generated using MEME v5.0.5. Transmembrane and cytosolic pentapeptides were predicted by the Phobius webserver. **B) VSP and** Ψ**VSP evolutionary dynamics**. The ΨVSP arrays (defined as >3 ΨVSP genes arrayed) in the *G. muris* genome arranged by chromosome, termini arranged to the right. Arrays of ΨVSPs without chromosomal context are shown below as orphan arrays. **C)** VSP cluster mostly together in a phylogeny whereas most ΨVSP gene arrays show relaxed clustering. Intact VSP genes are colored red, while ΨVSP genes are colored according to their resident ΨVSP array in panel A. Genes with black label are not found in arrays. **D) Expression measurements of VSP and** Ψ**VSP in *G. muris***. VSP and ΨVSP RNA-seq read counts (FPKM) from trophozoites, cysts and excyzoites. Size of the circle corresponds to expression level.

**Figure S6. A) Phylogenetic tree of *G. muris* and *G. intestinalis* NEKs. B) Phylogenetic tree of *G. muris* and *G. intestinalis* ARPs. C) Network analysis of *G. muris* and *G. intestinalis* NEKs and ARPs**. *G. muris* genes are represented in circle whereas *G. intestinalis* genes are shown in triangle. Blue indicates NEK and pink as ARP. Edges are weighted and scaled by reciprocal BLAST scores with evalue 1e-05 as the cutoff.

**Figure S7. Phylogenetic tree of C1 family cysteine proteases in diplomonads**. Black: *G. muris*, Red: *S. salmonicida*, Cyan: *G. intestinalis* GS/M, Green: *G. intestinalis* P15, Purple: *G. intestinalis* WB.

**Figure S8. Phylogenetic analyses of HGT genes. A)** Fructokinase. **B)** Mannose-6-phosphate isomerase. **C)** Transcriptional regulator/sugar kinase. **D)** Arginase. **E)** 2,5-diketo-D-gluconic acid reductase 1. **F)** 2,5-diketo-D-gluconic acid reductase 2. **G)** Carboxymuconolactone decarboxylase. **H)** Ketosteroid isomerase. **I)** Tryptophanase. **J)** Tae4. **K)** Quorum-quenching N-acyl-homoserine lactonase.

**Table S1. A)** One-to-one ortholog list between *G. muris* and *G. intestinalis* WB. Orthologs inferred by OrthoMCL. **B)** RNA-Seq expression values in excel file. Sorted according to FPKM values. **C)** VSP and ΨVSP gene metadata. ‘-’ in the pentapeptide motif represents lack of the 3’ end sequence. **D)** List of ventral disc proteins in *G. muris*. **E)** Proteases detected in *G. muris* and *G. intestinalis*. **F)** Detailed information about hits of unique genes in *G. muris*.

**Methods S1. Additional methods section**

## Acknowledgements

Elaine Hanson is acknowledged for technical support performing trophozoite purifications from infected mice. Phylogenetic analyses were performed on resources provided by the Swedish National Infrastructure for Computing (SNIC) at Uppsala Multidisciplinary Center for Advanced Computational Science (UPPMAX).

## Declarations

### Ethics approval and consent to participate

All animal studies were approved by the Institutional Animal Care and Use Committees of the University of California, San Diego, USA.

### Consent for publication

No applicable.

### Availability of data and material

Raw DNA and RNA sequence reads are archived at NCBI Sequence Read Archive (SRA) under accession number SRR8858297 - SRR8858305.

This Whole Genome Shotgun project has been deposited at DDBJ/EMBL/GenBank under the accession PRJNA524057. The version described in this paper is version VDLU00000000.1. The datasets generated and analyzed in this project are also available via GiardiaDB.

### Authors’ contributions

SGS, LE, JA and JJH conceived the study. JJH and EE performed RNA and DNA extractions. FX assembled the genome. FX, AJG, JJH, AA, DP, JA, SGS and EE annotated the genome. FX, JJH, AJG and SGS analyzed the data. All authors wrote the paper.

