## Supplementary figures and images for "The compact genome of *Giardia muris* reveals important steps in the evolution of intestinal protozoan parasites"

### Figure S1

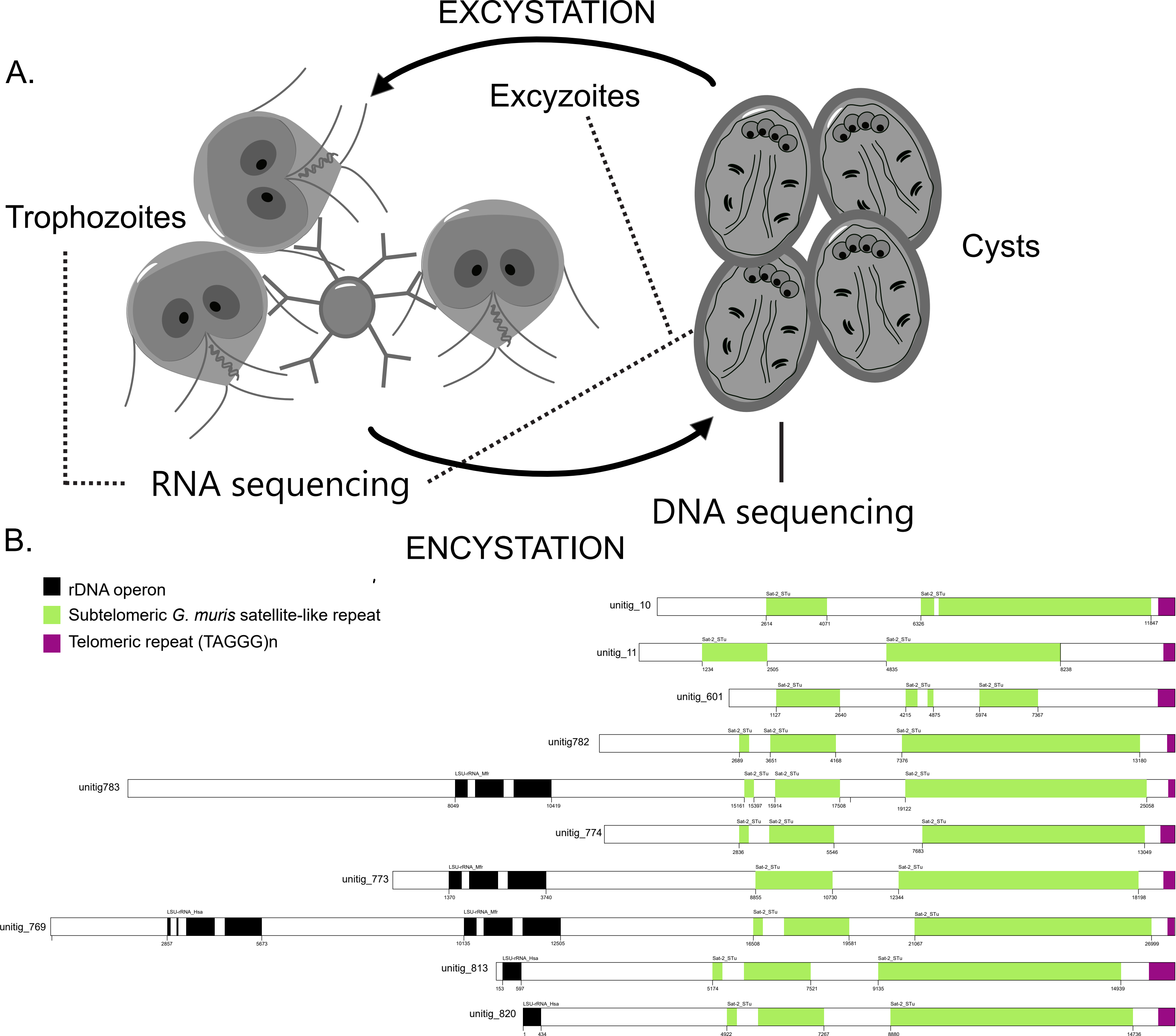

### Figure S2

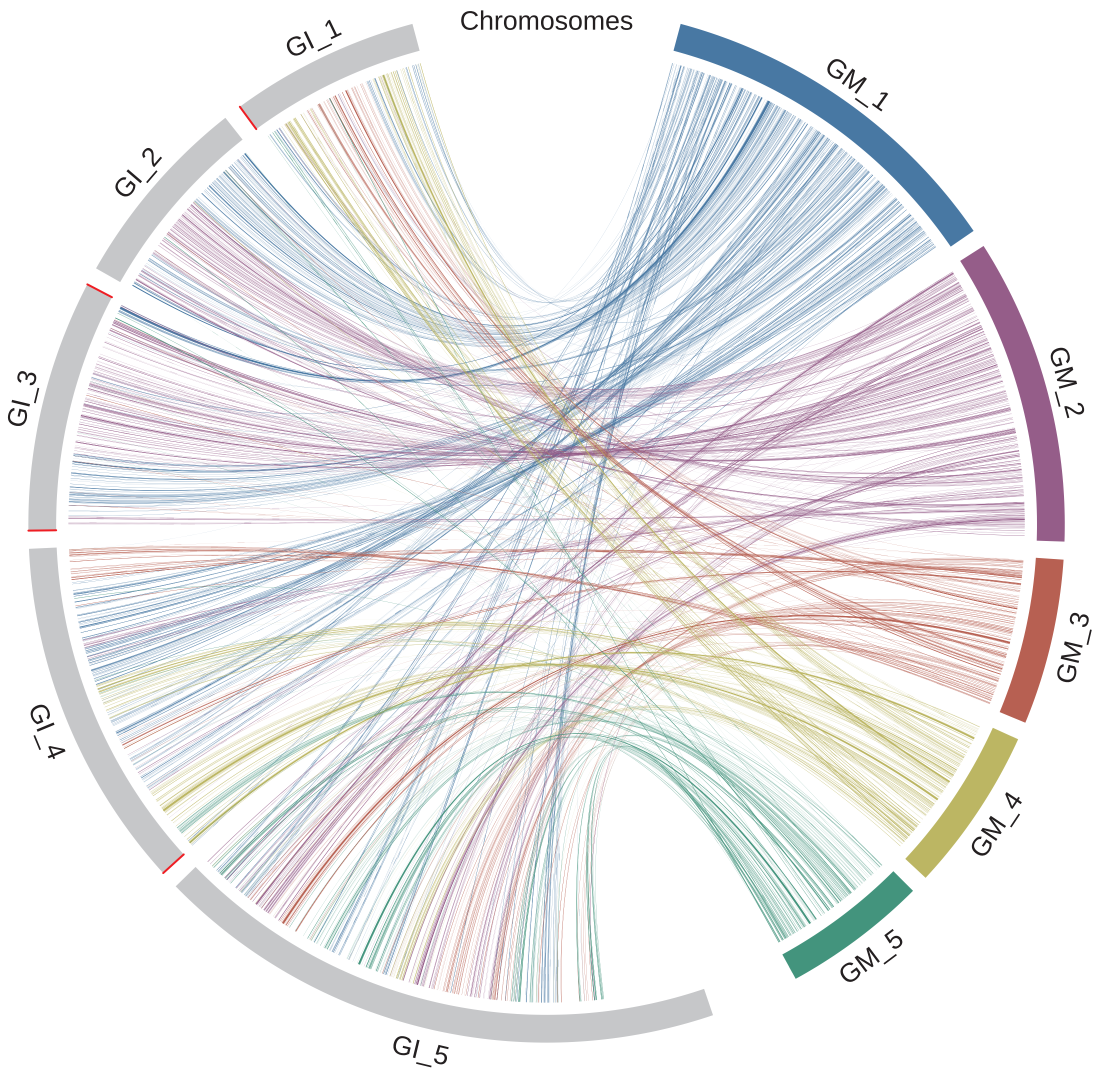

### Figure S5

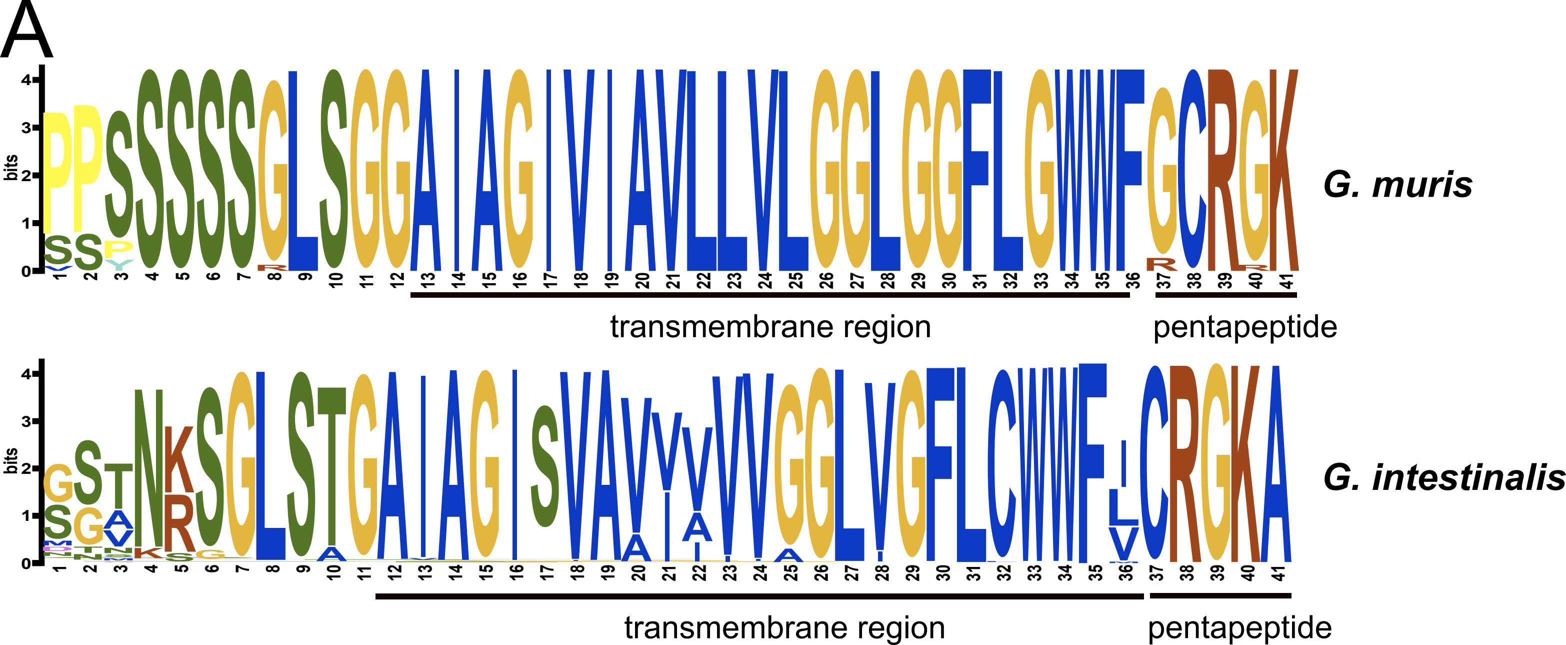

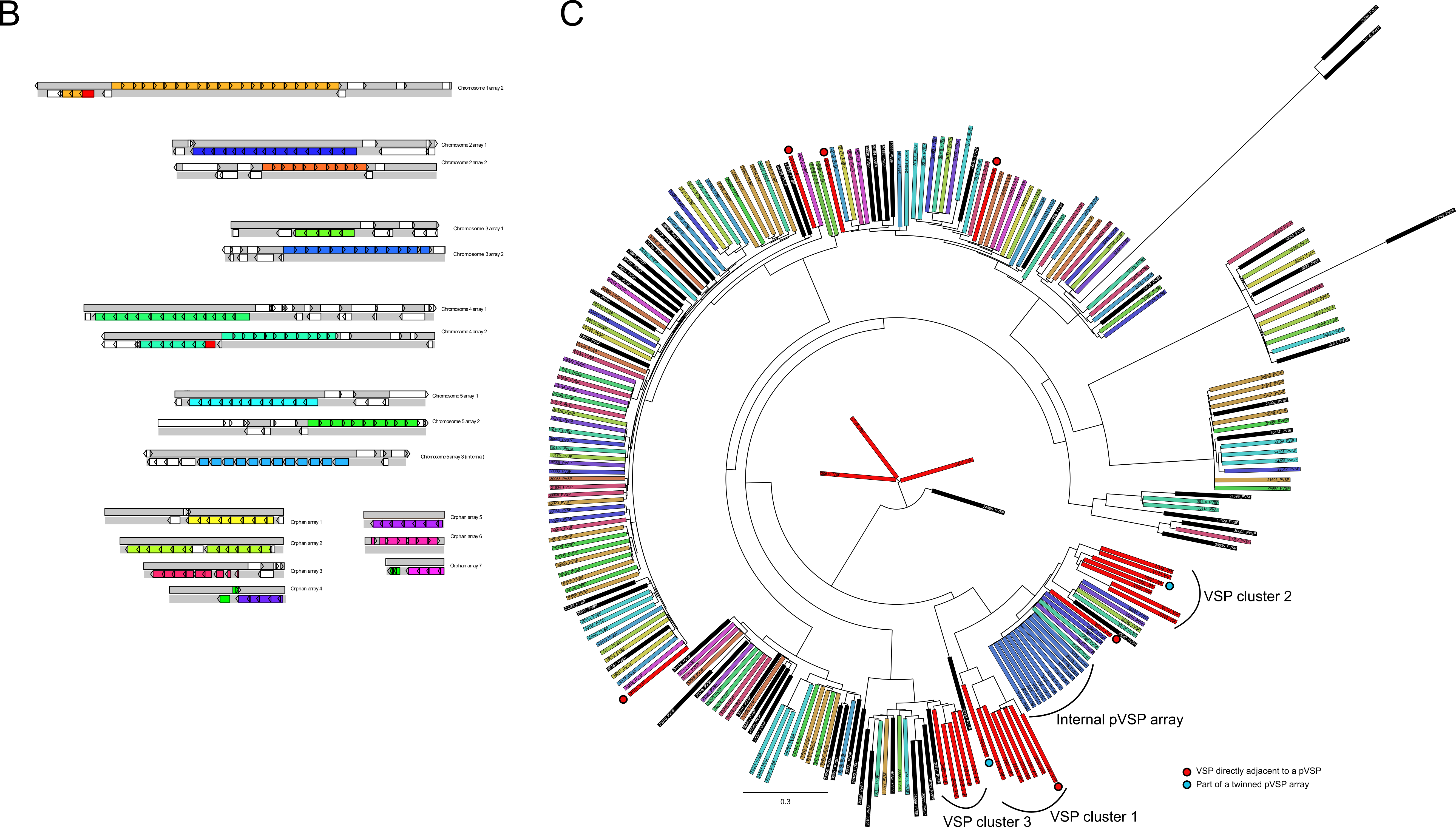

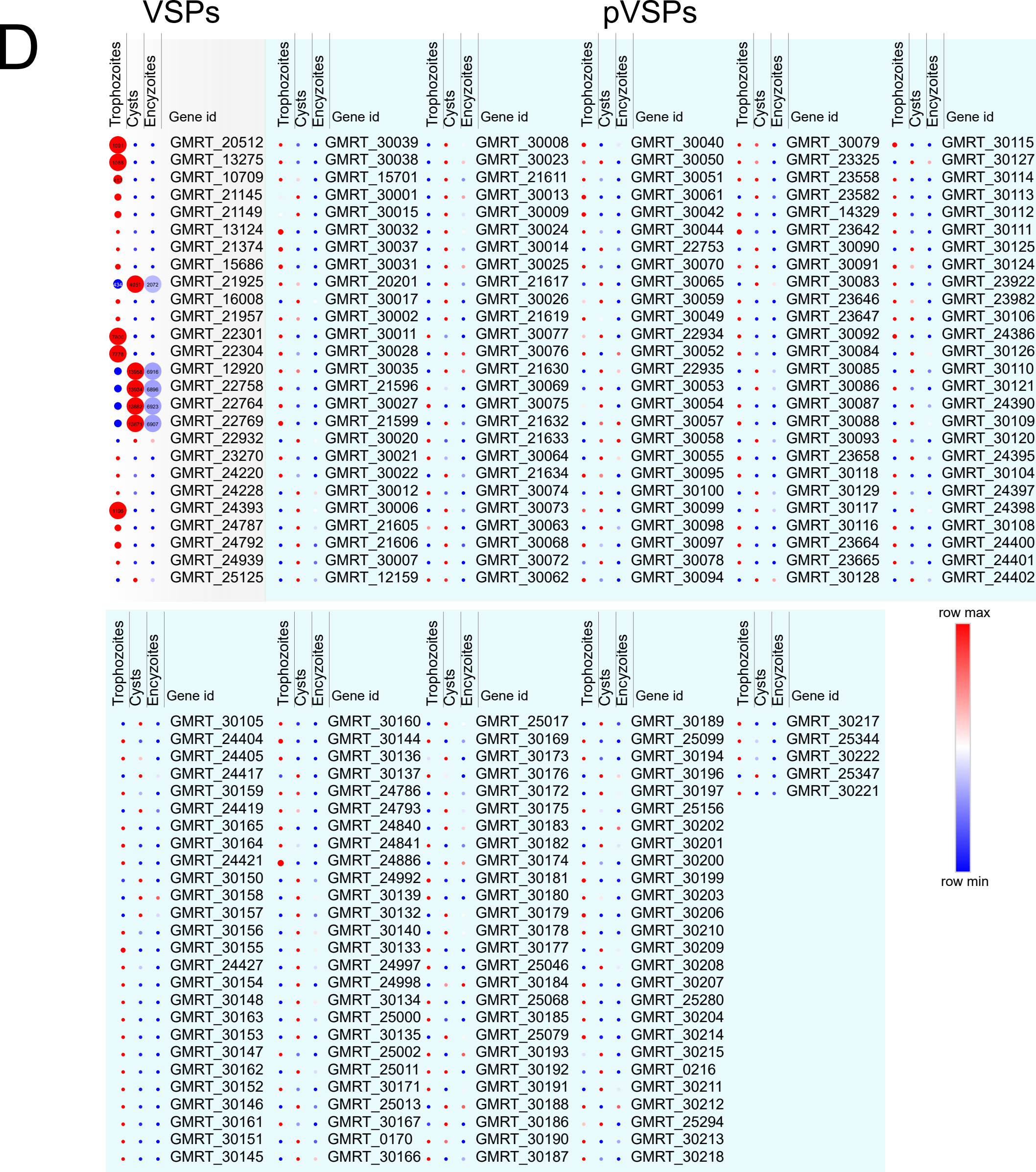

### Figure S6

A

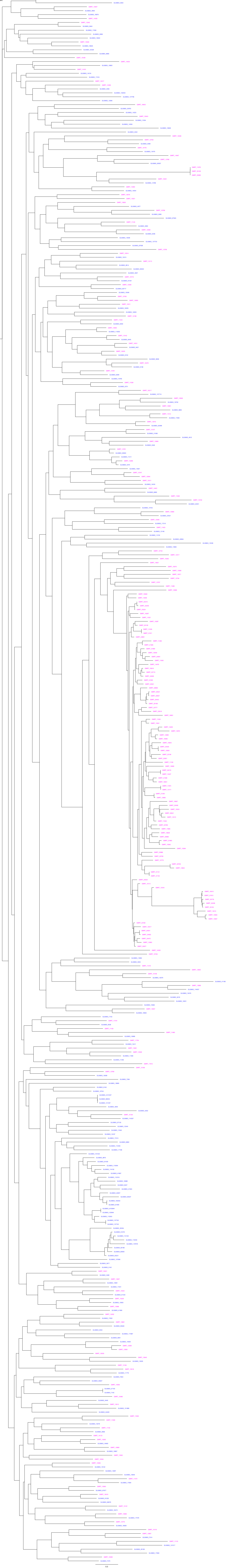

B

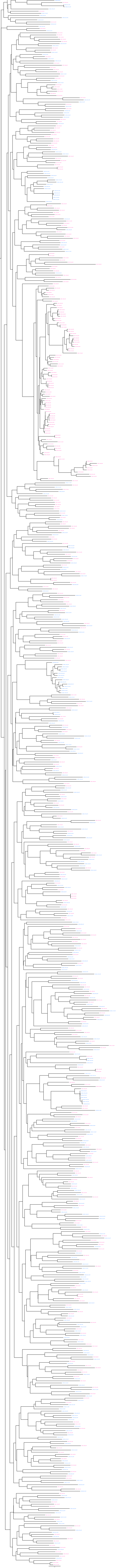

C

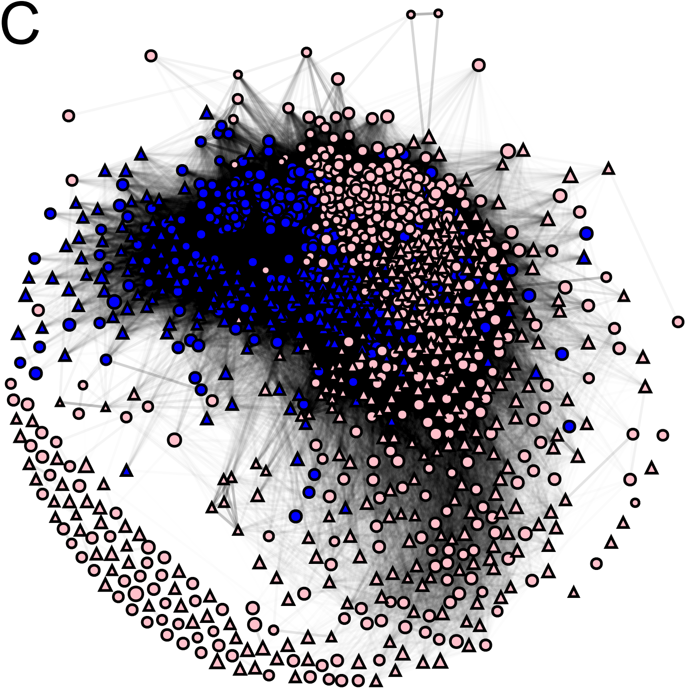

### Figure S7

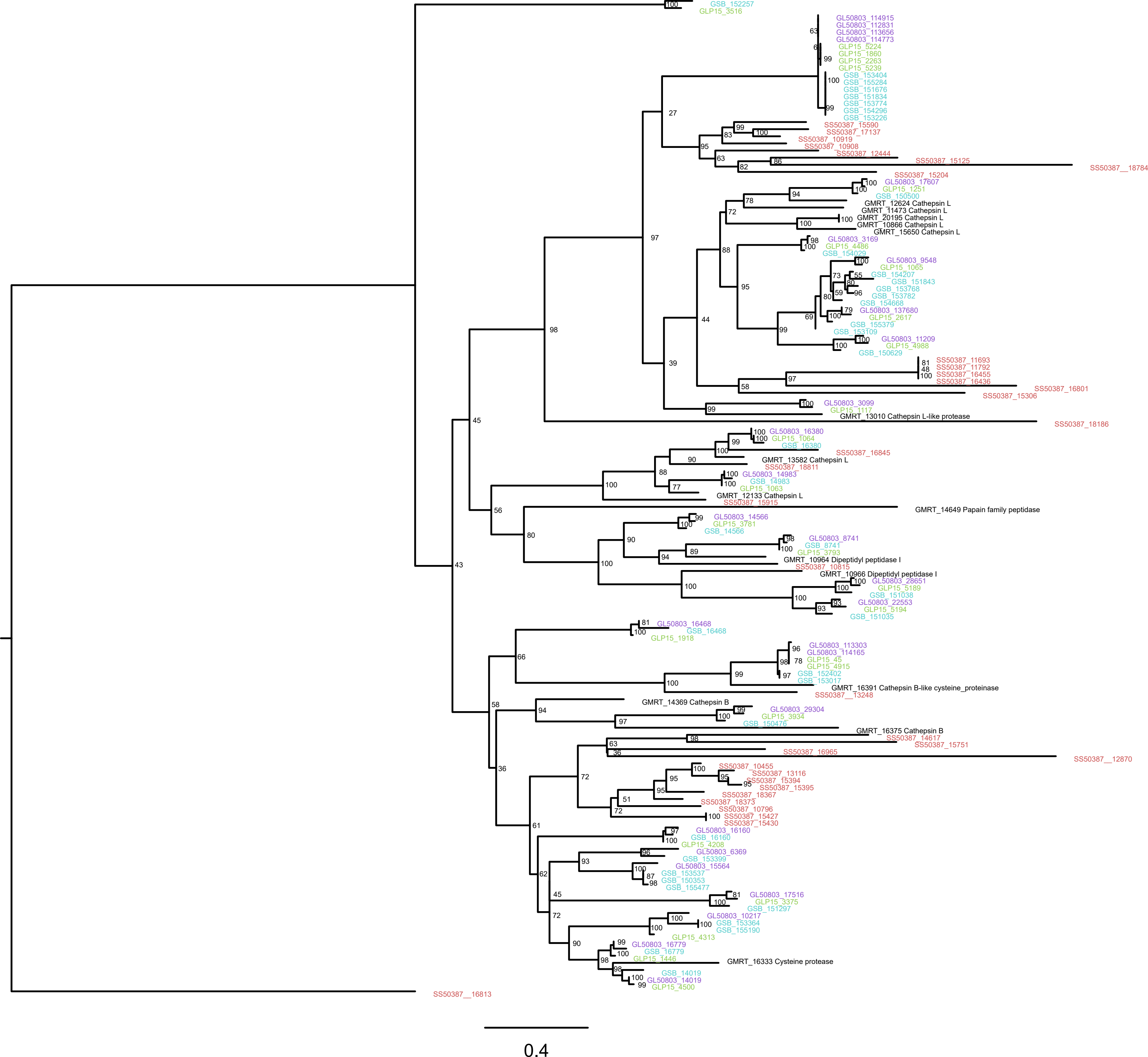
