## Supplementary material for "The compact genome of *Giardia muris* reveals important steps in the evolution of intestinal protozoan parasites": Figure S3

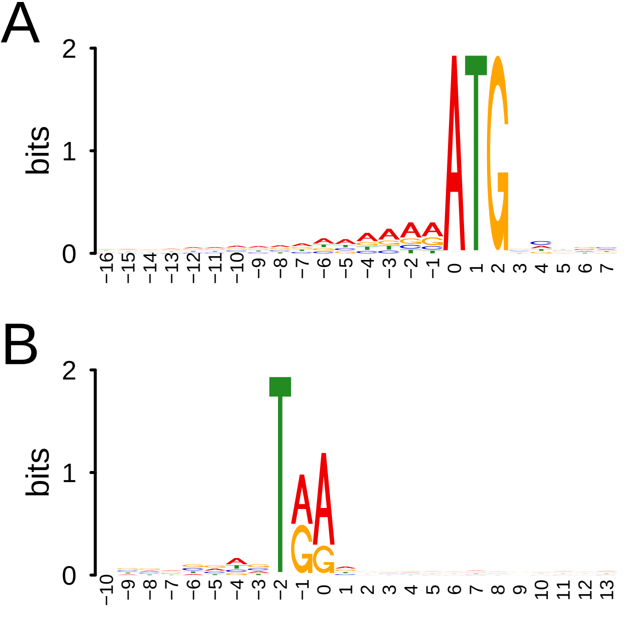
A
B
bits
0
1
2
−16 −15 −14 −13 −12 −11 −10 −9 −8 −7 −6 −5 −4 −3 −2 −1 0 1 2 3 4 5 6 7
−10 −9 −8 −7 −6 −5 −4 −3 −2 −1 0 1 2 3 4 5 6 7 8 9 10 11 12 13

C

| LOG2 Trophs/Cysts | Encyst up 90 min | Encyst up 7h | Encyst up 22h | Gene id |
|---|---|---|---|---|
| 9 | | | | GMRT_13982 |
| 9 | | ● | ● | GMRT_13204 |
| 9 | | ● | ● | GMRT_10774 |
| 9 | | | ● | GMRT_12407 |
| 8 | | | | GMRT_13942 |
| 8 | | | ● | GMRT_13341 |
| 8 | ● | ● | ● | GMRT_13400 |
| 8 | | ● | ● | GMRT_16219 |
| 8 | | | ● | GMRT_13851 |
| 8 | | | ● | GMRT_10589 |
| 8 | | | ● | GMRT_15202 |
| 8 | | | ● | GMRT_14302 |
| 7 | | | | GMRT_14494 |
| 7 | | ● | ● | GMRT_11770 |
| 7 | | | ● | GMRT_12514 |
| 7 | | | ● | GMRT_13776 |
| 7 | | | | GMRT_12207 |
| 7 | | ● | ● | GMRT_10157 |
| 7 | | | ● | GMRT_12519 |
| 7 | ● | | | GMRT_10224 |
| 7 | | | | GMRT_15543 |
| 7 | | | | GMRT_15329 |
| 7 | | ● | ● | GMRT_15984 |
| 7 | | | | GMRT_10478 |
| 7 | | | | GMRT_13283 |
| 7 | | | ● | GMRT_13422 |
| 7 | | | ● | GMRT_11433 |
| 6 | | | | GMRT_13315 |
| 6 | | | ● | GMRT_12667 |
| 6 | | | | GMRT_12476 |
| 6 | | | | GMRT_15303 |
| 6 | | | | GMRT_13901 |
| 6 | ● | ● | | GMRT_12279 |
| 6 | | | | GMRT_12528 |
| 6 | | | | GMRT_15601 |
| 6 | | | | GMRT_15514 |
| 6 | | | | GMRT_13605 |
| 6 | ● | | | GMRT_14361 |
| 6 | | | | GMRT_14093 |
| 6 | | | | GMRT_10233 |

| LOG2 Trophs/Cysts | Encyst up 90 min | Encyst up 7h | Encyst up 22h | Gene id |
|---|---|---|---|---|
| 6 | | ● | ● | GMRT_10574 |
| 6 | | | | GMRT_14982 |
| 6 | | | ● | GMRT_15986 |
| 6 | | | | GMRT_12453 |
| 6 | | | ● | GMRT_12189 |
| 6 | | | ● | GMRT_10312 |
| 6 | | | | GMRT_12190 |
| 6 | | | | GMRT_14841 |
| 6 | | | ● | GMRT_12243 |
| 6 | | | | GMRT_16336 |
| 6 | | | ● | GMRT_10833 |
| 6 | | ● | | GMRT_14208 |
| 6 | | | | GMRT_11693 |
| 6 | | ● | ● | GMRT_11117 |
| 6 | | | ● | GMRT_14265 |
| 6 | | | ● | GMRT_14586 |
| 6 | | | | GMRT_10367 |
| 6 | | | ● | GMRT_11957 |
| 6 | | | | GMRT_12759 |
| 6 | | | ● | GMRT_15885 |
| 6 | | | | GMRT_11905 |
| 6 | | | ● | GMRT_15547 |
| 6 | ● | | | GMRT_10756 |
| 6 | | | ● | GMRT_13994 |
| 5 | | | | GMRT_10377 |
| 5 | ● | | ● | GMRT_10123 |
| 5 | | | ● | GMRT_13780 |
| 5 | | ● | ● | GMRT_15811 |
| 5 | | | ● | GMRT_14189 |
| 5 | | | | GMRT_14883 |
| 5 | | | ● | GMRT_14997 |
| 5 | | | | GMRT_12394 |
| 5 | | | | GMRT_10848 |
| 5 | | | | GMRT_12419 |
| 5 | | | | GMRT_10135 |
| 5 | | | | GMRT_15872 |
| 5 | | | ● | GMRT_16332 |
| 5 | | | | GMRT_14187 |
| 5 | | | | GMRT_11145 |
| 5 | | | | GMRT_12776 |

| LOG2 Trophs/Cysts | Encyst up 90 min | Encyst up 7h | Encyst up 22h | Gene id |
|---|---|---|---|---|
| 5 | | | | GMRT_11843 |
| 5 | | | | GMRT_10541 |
| 5 | | | | GMRT_15517 |
| 5 | | | | GMRT_10471 |
| 5 | | | ● | GMRT_15257 |
| 5 | | | | GMRT_10501 |
| 5 | | | | GMRT_12607 |
| 5 | | | ● | GMRT_14295 |
| 5 | | | | GMRT_11315 |
| 5 | | | | GMRT_12559 |
| 5 | | | | GMRT_13415 |
| 5 | | | | GMRT_16213 |
| 5 | ● | | | GMRT_13109 |
| 5 | | | | GMRT_10567 |
| 5 | | | ● | GMRT_14575 |
| 5 | | | | GMRT_12715 |
| 5 | | | ● | GMRT_15985 |
| 5 | | | | GMRT_13841 |
| 5 | | | ● | GMRT_10038 |
| 5 | | | ● | GMRT_11441 |
| 5 | | | | GMRT_13609 |
| 5 | | | ● | GMRT_11004 |
| 5 | | ● | ● | GMRT_14079 |
| 5 | | | | GMRT_11698 |
| 5 | | ● | | GMRT_12381 |
| 5 | | | ● | GMRT_10030 |
| 5 | ● | | | GMRT_10585 |
| 5 | | | ● | GMRT_15240 |
| 5 | | | | GMRT_14576 |
| 5 | | | ● | GMRT_10991 |
| 5 | | | ● | GMRT_10607 |
| 5 | | | ● | GMRT_11520 |
| 5 | | | | GMRT_10127 |
| 5 | | | | GMRT_15829 |
| 5 | | ● | ● | GMRT_14224 |
| 5 | ● | ● | ● | GMRT_13401 |
| 5 | | | | GMRT_12332 |
| 5 | | | | GMRT_13227 |
| 5 | | | | GMRT_14385 |
| 5 | | | ● | GMRT_10590 |

| LOG2 Trophs/Cysts | Encyst up 90 min | Encyst up 7h | Encyst up 22h | Gene id |
|---|---|---|---|---|
| 5 | | | ● | GMRT_10989 |
| 5 | | | | GMRT_12531 |
| 5 | | ● | ● | GMRT_11758 |
| 5 | | | | GMRT_15162 |
| 5 | | | | GMRT_10361 |
| 5 | | | | GMRT_13783 |
| 5 | ● | | | GMRT_13387 |
| 5 | | | ● | GMRT_14264 |
| 5 | | | ● | GMRT_12874 |
| 5 | | | ● | GMRT_11847 |
| 4 | | | | GMRT_13488 |
| 4 | | | | GMRT_13010 |
| 4 | | | ● | GMRT_16375 |
| 4 | | | | GMRT_14214 |
| 4 | | | | GMRT_12291 |
| 4 | | | | GMRT_12942 |
| 4 | | | ● | GMRT_13845 |
| 4 | | | ● | GMRT_13363 |
| 4 | | | | GMRT_10998 |
| 4 | | | | GMRT_13929 |
| 4 | | | | GMRT_16145 |
| 4 | | | | GMRT_16374 |
| 4 | | | | GMRT_12385 |
| 4 | | | ● | GMRT_14301 |
| 4 | | | | GMRT_14584 |
| 4 | | | ● | GMRT_11013 |
| 4 | | | ● | GMRT_10770 |
| 4 | | | ● | GMRT_13180 |
| 4 | ● | | | GMRT_15317 |
| 4 | | | | GMRT_10811 |
| 4 | | | | GMRT_12821 |
| 4 | | | | GMRT_12091 |
| 4 | | | | GMRT_11966 |
| 4 | | | ● | GMRT_12130 |
| 4 | | | | GMRT_13902 |
| 4 | | | | GMRT_16379 |
| 4 | | | | GMRT_14952 |
| 4 | | | | GMRT_13245 |
| 4 | ● | | | GMRT_15524 |
| 4 | | | | GMRT_15258 |

| LOG2 Trophs/Cysts | Encyst up 90 min | Encyst up 7h | Encyst up 22h | Gene id |
|---|---|---|---|---|
| 4 | | | | GMRT_12300 |
| 4 | | | | GMRT_16311 |
| 4 | | | | GMRT_15163 |
| 4 | | | ● | GMRT_12285 |
| 4 | | | | GMRT_14430 |
| 4 | | | ● | GMRT_13601 |
| 4 | ● | ● | | GMRT_12672 |
| 4 | | | ● | GMRT_15899 |
| 4 | | | ● | GMRT_15546 |
| 4 | ● | | | GMRT_13611 |
| 4 | | | ● | GMRT_14303 |
| 4 | | | | GMRT_10495 |
| 4 | | | | GMRT_15159 |
| 4 | | ● | | GMRT_10188 |
| 4 | | | | GMRT_11987 |
| 4 | | | | GMRT_14267 |
| 4 | ● | | | GMRT_13101 |
| 4 | | | | GMRT_16305 |
| 4 | | | | GMRT_12824 |
| 4 | | | | GMRT_10401 |
| 4 | ● | | ● | GMRT_14423 |
| 4 | | | | GMRT_10296 |
| 4 | ● | ● | ● | GMRT_11676 |
| 4 | | | | GMRT_13096 |
| 4 | | | ● | GMRT_13390 |
| 4 | | | ● | GMRT_15222 |
| 4 | | | | GMRT_13985 |
| 4 | | | | GMRT_12620 |
| 4 | ● | | | GMRT_10467 |
| 4 | | ● | ● | GMRT_15068 |
| 4 | | | | GMRT_12343 |
| 4 | | ● | ● | GMRT_13794 |
| 4 | | | ● | GMRT_12815 |
| 4 | | ● | | GMRT_13570 |
| 4 | | | | GMRT_11953 |
| 4 | | | | GMRT_14849 |
| 4 | | | | GMRT_12799 |
| 4 | | | | GMRT_13282 |
| 4 | | | | GMRT_10492 |
