## Supplementary material for "The compact genome of *Giardia muris* reveals important steps in the evolution of intestinal protozoan parasites": Figure S4

A.

*Trans-spliced introns*

```
GMRT_12388 GTgtatgtgtg.../ /...ggactaacacgcagCT
GMRT_11576 AGgtatgtccg.../ /...tgactaacacgcagCA
GMRT_10332 GGgtatgtggt.../ /...ttactaacacgcagCT
GMRT_11888 TGatatgttgt.../ /...tcactaacacgcagAC
GMRT_22684 TGgtatggccc.../ /...ttactaacacgcagTC
             .****..             ************
```

*Cis-spliced introns*

```
GMRT_fx013 GCgtatgtatc    tc    catctaacacgtagAA
GMRT_15227 GGgtatgtgcc (124nt) tcactaacgcgcagAT
GMRT_11066 CCgtatgttca  tatcg  tctctaacacgcagCT
             ******.             .*****.**.**
```

B.

```
GMRT_12388-1     gtatgtgtgtgcgtcagccgctgtgtgttgtgtgtttcctttactcaatgacggtctc

GMRT_12388-1d    gtatgtgtgtgcgtcagccgctgtgtgttgtgtgtttcctttacttaaatggttc

GMRT_12388-2     aaaaatcagcggctgacgcgcaggaccatctggactaacacgcag


GMRT_11576-1     gtatgtccgctgtgtgcgagcatgtgtgttgtgtgtctcctttactcaacttccgcgg

GMRT_11576-1d    gtatgtccgctgtgtgcgagcatgtgtgttgtgtgtctcctttactcaatatcaaattctccattgtatttcat

GMRT_11576-2     aaaaaacctcaagtactggagggaagaaaagctcgcaccagcagaatcttgactaacacgcag

GMRT_11576-2-gt gtatgtggtgttccgcttgctgtgtgttgtgtgttccttcacttaagttctgtc

GMRT_11576-3     aaaaagagacacgaggacgatcggcaagcggaacacacagtcctttactaacacgcag


GMRT_11888-1     atatgttgtgcgtttcaccccgttgtgtgttgtgtgttccttttctcaagcattggct

GMRT_11888-2     aaaacggggtgaacgtgcagtcctcactaacacgcag


GMRT_22684-1     gtatggcccctcctccttcactcaagccttct

GMRT_22684-2     aaaggagggaccaatcttactaacacgcag
```

C.

**Cleavage motif**

```
tcctttactcaa
tcctttacttaa
tcctttactcaa
tccttcacttaa
tccttttctcaa
tccttcactcaa
*****..**.**
```

D.

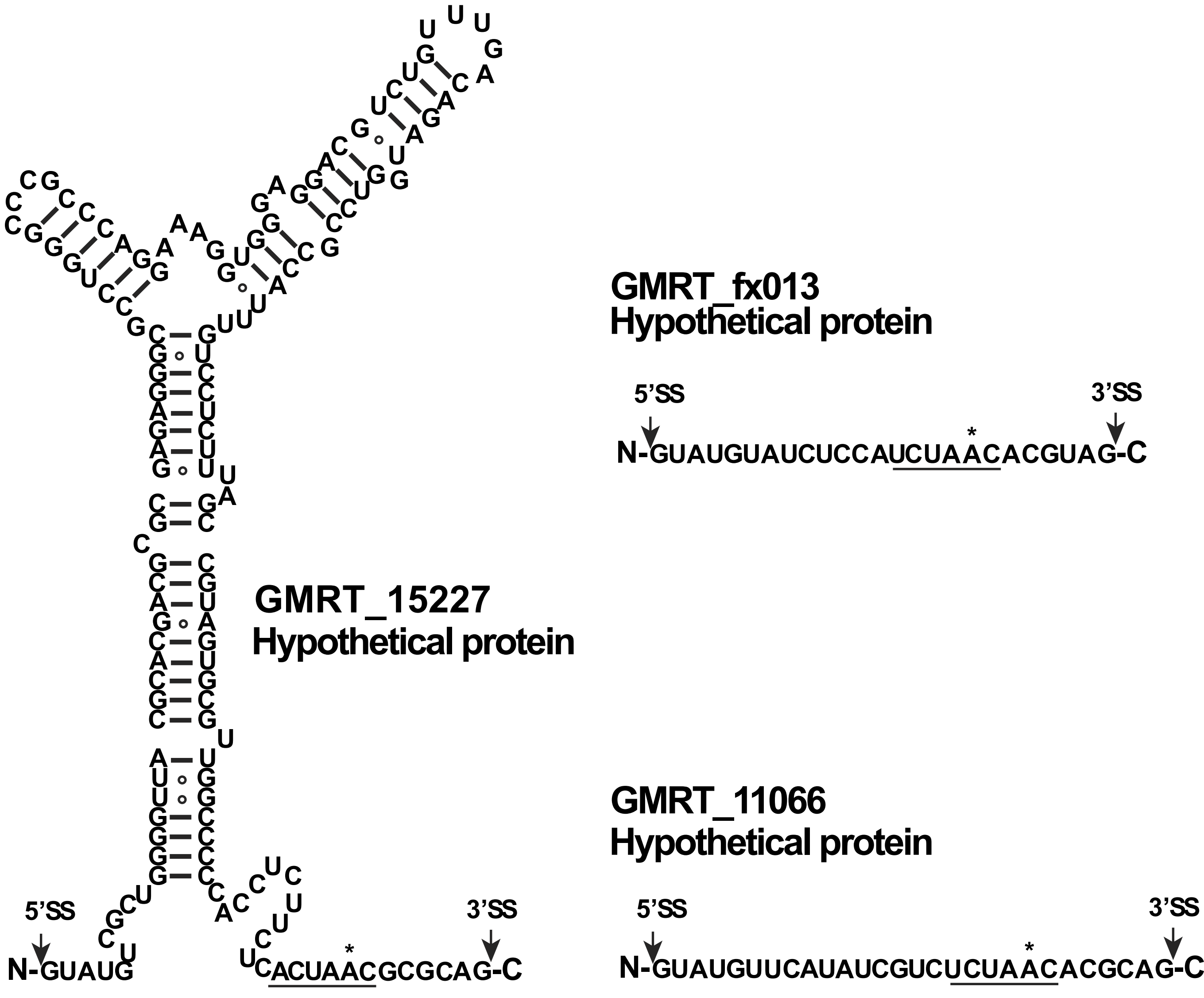
GMRT_fx013
Hypothetical protein
5’SS
3’SS
N-GUAUGUAUCUCCAUCUAACACGUAG-C
GMRT_15227
Hypothetical protein
5’SS
3’SS
N-GUAUG
UCACUAACGCGCAG-C
GMRT_11066
Hypothetical protein
5’SS
3’SS
N-GUAUGUUCAUAUCGUCUCUAACACGCAG-C

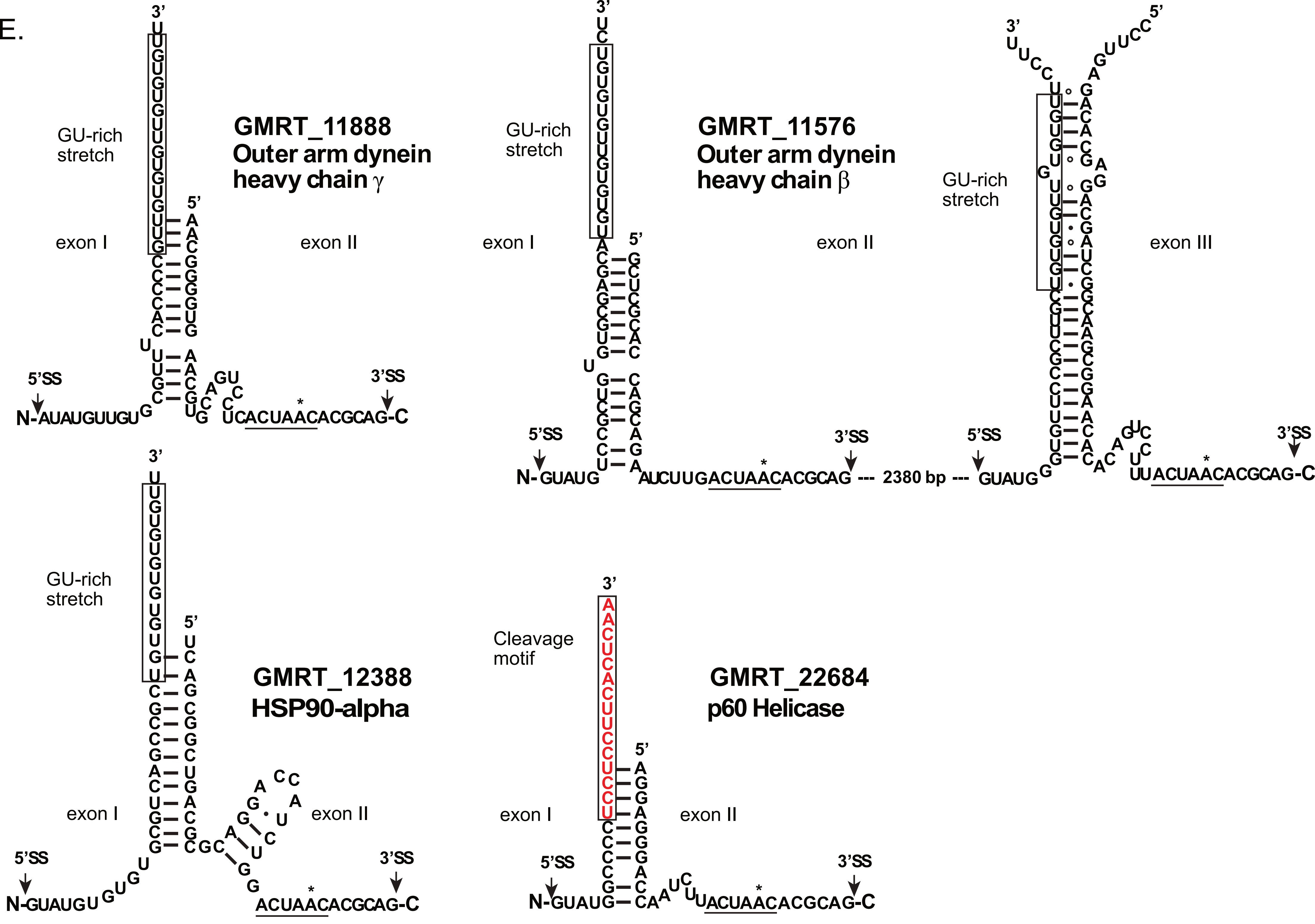

E.
GMRT_11888
Outer arm dynein heavy chain γ
GU-rich stretch
exon I
exon II
5'SS
3'SS
N-AUAUGUUGU
UCACUAACACGCAG-C
GMRT_11576
Outer arm dynein heavy chain β
GU-rich stretch
exon I
exon II
exon III
5'SS
3'SS
N-GUAUG
AUCUUGACUAACACGCAG --- 2380 bp --- GUAUG
UUACUAACACGCAG-C
GMRT_12388
HSP90-alpha
GU-rich stretch
exon I
exon II
5'SS
3'SS
N-GUAUGU
ACUAACACGCAG-C
GMRT_22684
p60 Helicase
Cleavage motif
exon I
exon II
5'SS
3'SS
N-GUAUG
UACUAACACGCAG-C
