## Supplementary material for "The compact genome of *Giardia muris* reveals important steps in the evolution of intestinal protozoan parasites": Figure S8

### A. Fructokinase

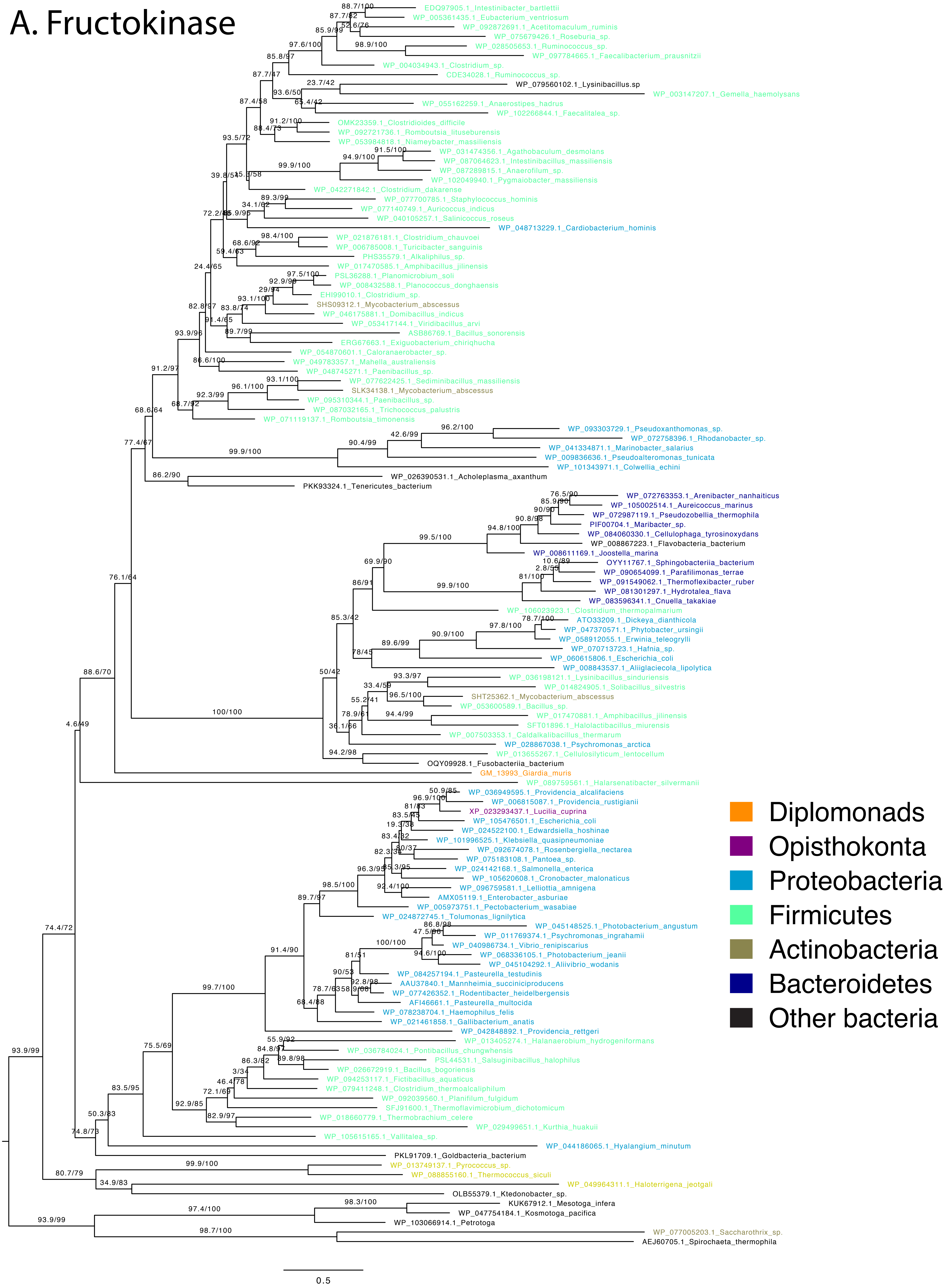

B. ManA

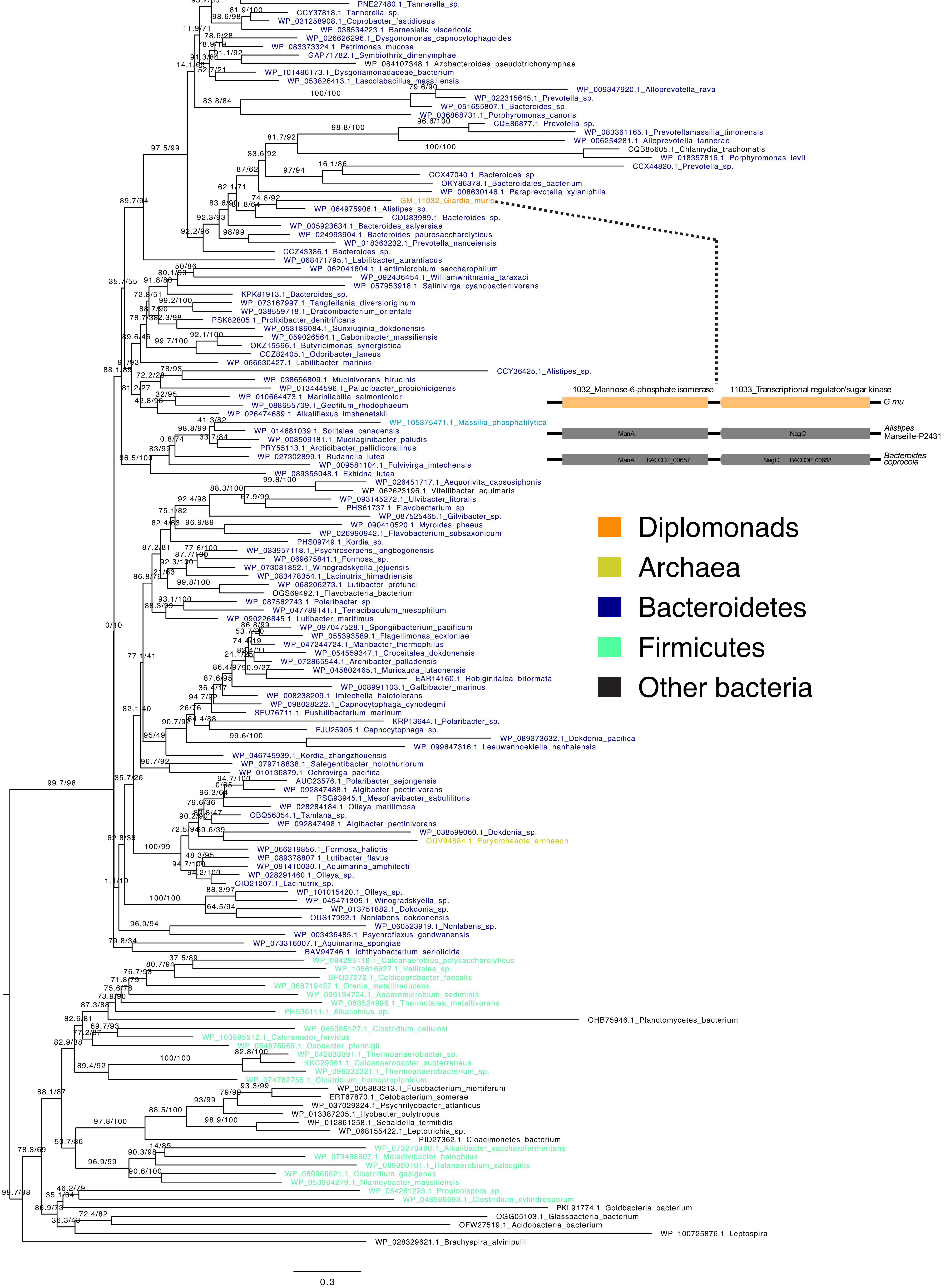
WP_080905946.1_Parabacteroides_sp.
WP_036880967.1_Porphyromonas_sp.
WP_087330492.1_Parabacteroides_sp.
WP_085536262.1_Massilibacteroides_vaginae
PNE27480.1_Tannerella_sp.
CCY37818.1_Tannerella_sp.
WP_031258908.1_Coprobacter_fastidiosus
WP_038534223.1_Barnesiella_viscericola
WP_026626296.1_Dysgonomonas_capnocytophagoides
WP_083373324.1_Petrimonas_mucosa
GAP71782.1_Symbiothrix_dinenymphae
WP_084107348.1_Azobacteroides_pseudotrichonymphae
WP_101486173.1_Dysgonamonadaceae_bacterium
WP_053826413.1_Lascolabacillus_massiliensis
WP_009347920.1_Alloprevotella_rava
WP_022315645.1_Prevotella_sp.
WP_051655807.1_Bacteroides_sp.
WP_036868731.1_Porphyromonas_canoris
CDE86877.1_Prevotella_sp.
WP_083361165.1_Prevotellamassilia_timonensis
WP_006254281.1_Alloprevotella_tannerae
CQB85605.1_Chlamydia_trachomatis
WP_018357816.1_Porphyromonas_levii
CCX44820.1_Prevotella_sp.
CCX47040.1_Bacteroides_sp.
OKY86378.1_Bacteroidales_bacterium
WP_008630146.1_Paraprevotella_xylaniphila
GM_11032_Giardia_muris
WP_064975906.1_Alistipes_sp.
CDD83989.1_Bacteroides_sp.
WP_005923634.1_Bacteroides_salyersiae
WP_024993904.1_Bacteroides_paurosaccharolyticus
WP_018363232.1_Prevotella_nanceiensis
CCZ43386.1_Bacteroides_sp.
WP_068471795.1_Labilibacter_aurantiacus
WP_062041604.1_Lentimicrobium_saccharophilum
WP_092436454.1_Williamwhitmania_taraxaci
WP_057953918.1_Salinivirga_cyanobacteriivorans
KPK81913.1_Bacteroides_sp.
WP_073167997.1_Tangfeifania_diversioriginum
WP_038559718.1_Draconibacterium_orientale
PSK82805.1_Prolixibacter_denitrificans
WP_053186084.1_Sunxiuqinia_dokdonensis
WP_059026564.1_Gabonibacter_massiliensis
OKZ15566.1_Butyricimonas_synergistica
CCZ82405.1_Odoribacter_laneus
WP_066630427.1_Labilibacter_marinus
CCY36425.1_Alistipes_sp.
WP_038656809.1_Mucinivorans_hirudinis
WP_013444596.1_Paludibacter_propionicigenes
WP_010664473.1_Marinilabilia_salmonicolor
WP_088655709.1_Geofilum_rhodophaeum
WP_026474689.1_Alkaliflexus_imshenetskii
WP_105375471.1_Massilia_phosphatilytica
WP_014681039.1_Solitalea_canadensis
WP_008509181.1_Mucilaginibacter_paludis
PRY55113.1_Arcticibacter_pallidicorallinus
WP_027302899.1_Rudanella_lutea
WP_009581104.1_Fulvivirga_imtechensis
WP_089355048.1_Ekhidna_lutea
WP_026451717.1_Aequorivita_capsosiphonis
WP_062623196.1_Vitellibacter_aquimaris
WP_093145272.1_Ulvibacter_litoralis
PHS61737.1_Flavobacterium_sp.
WP_087525465.1_Gilvibacter_sp.
WP_090410520.1_Myroides_phaeus
WP_026990942.1_Flavobacterium_subsaxonicum
PHS09749.1_Kordia_sp.
WP_033957118.1_Psychroserpens_jangbogonensis
WP_069675841.1_Formosa_sp.
WP_073081852.1_Winogradskyella_jejuensis
WP_083478354.1_Lacinutrix_himadriensis
WP_068206273.1_Lutibacter_profundi
OGS69492.1_Flavobacteria_bacterium
WP_087562743.1_Polaribacter_sp.
WP_047789141.1_Tenacibaculum_mesophilum
WP_090226845.1_Lutibacter_maritimus
WP_097047528.1_Spongiibacterium_pacificum
WP_055393589.1_Flagellimonas_eckloniae
WP_047244724.1_Maribacter_thermophilus
WP_054559347.1_Croceitalea_dokdonensis
WP_072865544.1_Arenibacter_palladensis
WP_045802465.1_Muricauda_lutaonensis
EAR14160.1_Robiginitalea_biformata
WP_008991103.1_Galbibacter_marinus
WP_008238209.1_Imtechella_halotolerans
WP_098028222.1_Capnocytophaga_cynodegmi
SFU76711.1_Pustulibacterium_marinum
KRP13644.1_Polaribacter_sp.
EJU25905.1_Capnocytophaga_sp.
WP_089373632.1_Dokdonia_pacifica
WP_099647316.1_Leeuwenhoekiella_nanhaiensis
WP_046745939.1_Kordia_zhangzhouensis
WP_079718838.1_Salegentibacter_holothuriorum
WP_010136879.1_Ochrovirga_pacifica
AUC23576.1_Polaribacter_sejongensis
WP_092847488.1_Algibacter_pectinivorans
PSG93945.1_Mesoflavibacter_sabulilitoris
WP_028284184.1_Olleya_marilimosa
OBQ56354.1_Tamlana_sp.
WP_092847498.1_Algibacter_pectinivorans
WP_038599060.1_Dokdonia_sp.
OUV94884.1_Euryarchaeota_archaeon
WP_066219856.1_Formosa_haliotis
WP_089378807.1_Lutibacter_flavus
WP_091410030.1_Aquimarina_amphilecti
WP_028291460.1_Olleya_sp.
OIQ21207.1_Lacinutrix_sp.
WP_101015420.1_Olleya_sp.
WP_045471305.1_Winogradskyella_sp.
WP_013751882.1_Dokdonia_sp.
OUS17992.1_Nonlabens_dokdonensis
WP_060523919.1_Nonlabens_sp.
WP_003436485.1_Psychroflexus_gondwanensis
WP_073316007.1_Aquimarina_spongiae
BAV94746.1_Ichthyobacterium_seriolicida
WP_084295118.1_Caldanaerobius_polysaccharolyticus
WP_105616627.1_Vallitalea_sp.
SFQ27272.1_Caldicoprobacter_faecalis
WP_068719437.1_Orenia_metallireducens
WP_095134704.1_Anaeromicrobium_sediminis
WP_083524996.1_Thermotalea_metallivorans
PHS36111.1_Alkaliphilus_sp.
OHB75946.1_Planctomycetes_bacterium
WP_045085127.1_Clostridium_cellulosi
WP_103895512.1_Caloramator_fervidus
WP_054876969.1_Oxobacter_pfennigii
WP_042833391.1_Thermoanaerobacter_sp.
KKC29361.1_Caldanaerobacter_subterraneus
WP_096232321.1_Thermoanaerobacterium_sp.
WP_074782755.1_Clostridium_homopropionicum
WP_005883213.1_Fusobacterium_mortiferum
ERT67870.1_Cetobacterium_somerae
WP_037029324.1_Psychrilyobacter_atlanticus
WP_013387205.1_Ilyobacter_polytropus
WP_012861258.1_Sebaldella_termitidis
WP_068155422.1_Leptotrichia_sp.
PID27362.1_Cloacimonetes_bacterium
WP_073270490.1_Alkalibacter_saccharofermentans
WP_079488607.1_Maledivibacter_halophilus
WP_089860101.1_Halanaerobium_salsuginis
WP_089966621.1_Clostridium_gasigenes
WP_053984279.1_Niameybacter_massiliensis
WP_054261223.1_Propionispora_sp.
WP_048569693.1_Clostridium_cylindrosporum
PKL91774.1_Goldbacteria_bacterium
OGG05103.1_Glassbacteria_bacterium
OFW27519.1_Acidobacteria_bacterium
WP_100725876.1_Leptospira
WP_028329621.1_Brachyspira_alvinipulli
0.3
1032_Mannose-6-phosphate isomerase
11033_Transcriptional regulator/sugar kinase
G. mu
ManA
NagC
Alistipes Marseille-P2431
ManA BACCOP_00657
NagC BACCOP_00658
Bacteroides coprocola
Diplomonads
Archaea
Bacteroidetes
Firmicutes
Other bacteria

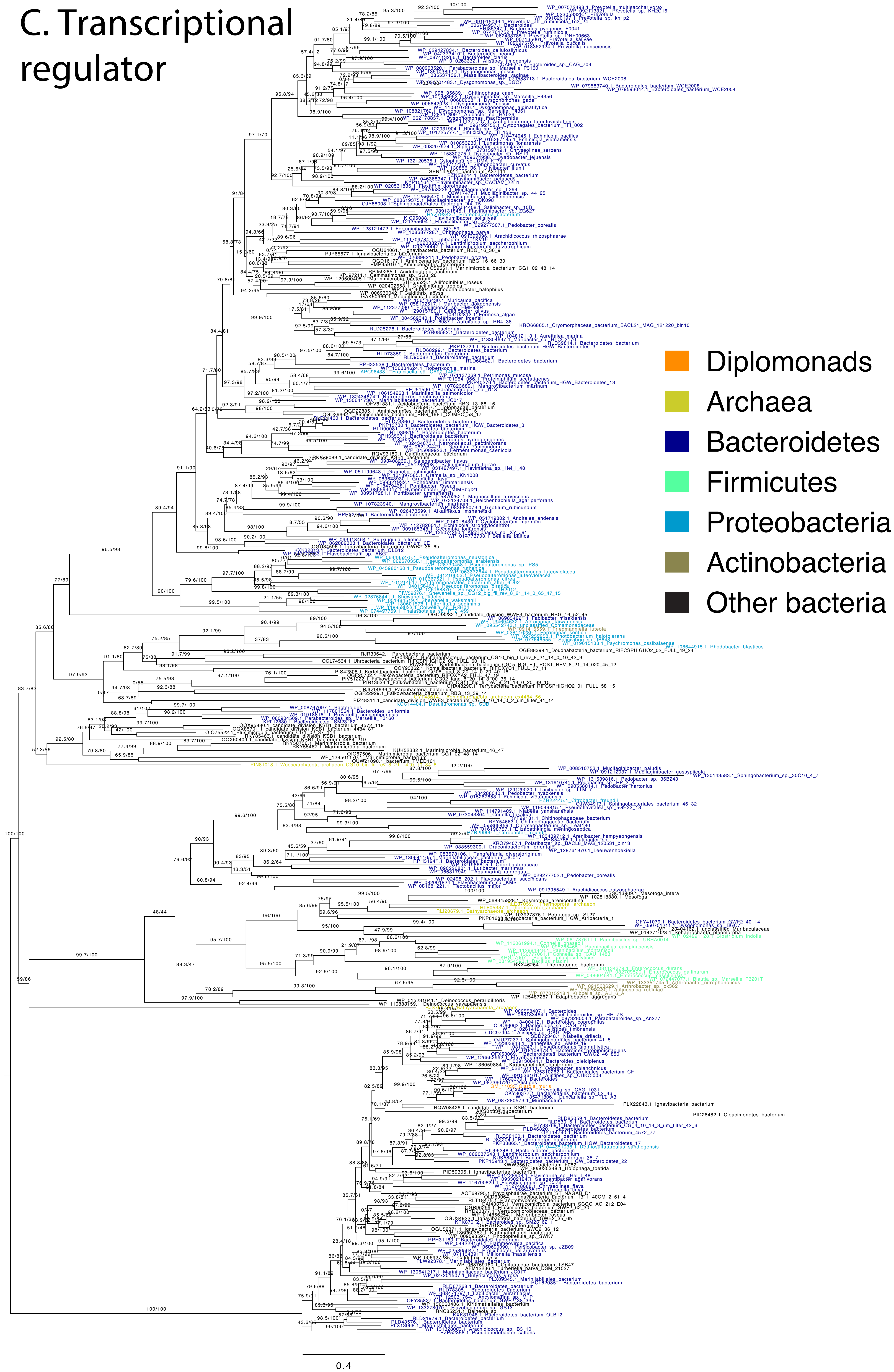
C. Transcriptional regulator
Diplomonads
Archaea
Bacteroidetes
Firmicutes
Proteobacteria
Actinobacteria
Other bacteria
0.4

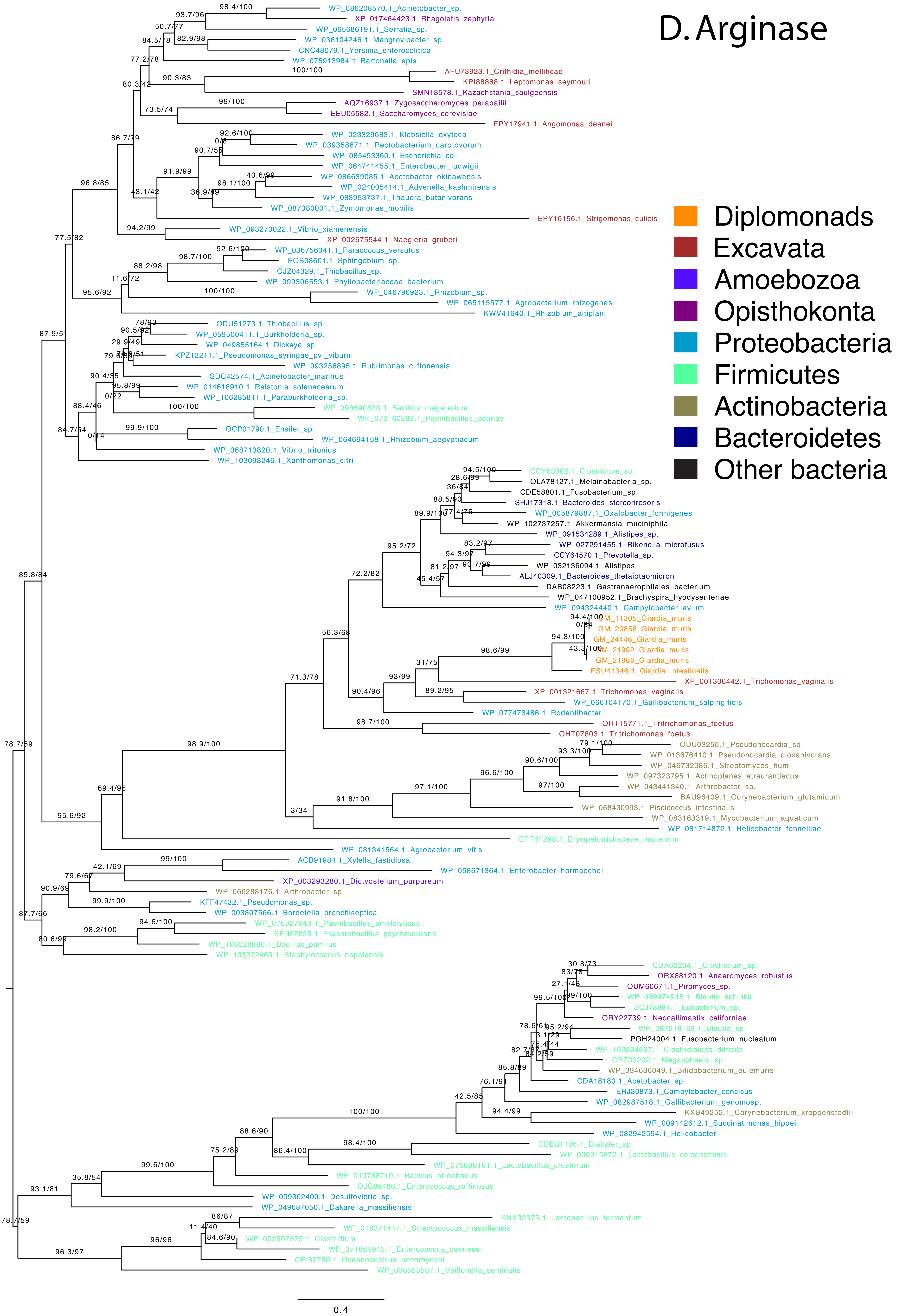
D. Arginase
Diplomonads
Excavata
Amoebozoa
Opisthokonta
Proteobacteria
Firmicutes
Actinobacteria
Bacteroidetes
Other bacteria
WP_086208570.1_Acinetobacter_sp.
XP_017464423.1_Rhagoletis_zephyria
WP_065686191.1_Serratia_sp.
WP_036104246.1_Mangrovibacter_sp.
CNC48079.1_Yersinia_enterocolitica
WP_075913984.1_Bartonella_apis
AFU73923.1_Crithidia_mellificae
KPI88868.1_Leptomonas_seymouri
SMN18578.1_Kazachstania_saulgeensis
AQZ16937.1_Zygosaccharomyces_parabailii
EEU05582.1_Saccharomyces_cerevisiae
EPY17941.1_Angomonas_deanei
WP_023329683.1_Klebsiella_oxytoca
WP_039358671.1_Pectobacterium_carotovorum
WP_085453360.1_Escherichia_coli
WP_064741455.1_Enterobacter_ludwigii
WP_086639085.1_Acetobacter_okinawensis
WP_024005414.1_Advenella_kashmirensis
WP_083953737.1_Thauera_butanivorans
WP_087380001.1_Zymomonas_mobilis
EPY16156.1_Strigomonas_culicis
WP_093270022.1_Vibrio_xiamenensis
XP_002675544.1_Naegleria_gruberi
WP_036756041.1_Paracoccus_versutus
EQB08601.1_Sphingobium_sp.
OJZ04329.1_Thiobacillus_sp.
WP_099306553.1_Phyllobacteriaceae_bacterium
WP_046796923.1_Rhizobium_sp.
WP_065115577.1_Agrobacterium_rhizogenes
KWV41640.1_Rhizobium_altiplani
ODU51273.1_Thiobacillus_sp.
WP_059500411.1_Burkholderia_sp.
WP_049855164.1_Dickeya_sp.
KPZ13211.1_Pseudomonas_syringae_pv._viburni
WP_093256895.1_Rubrimonas_cliftonensis
SDC42574.1_Acinetobacter_marinus
WP_014618910.1_Ralstonia_solanacearum
WP_106285811.1_Paraburkholderia_sp.
WP_098590838.1_Bacillus_megaterium
WP_076160383.1_Paenibacillus_peoriae
OCP01790.1_Ensifer_sp.
WP_064694158.1_Rhizobium_aegyptiacum
WP_068713820.1_Vibrio_tritonius
WP_103093246.1_Xanthomonas_citri
CCY63062.1_Clostridium_sp.
OLA78127.1_Melainabacteria_sp.
CDE58801.1_Fusobacterium_sp.
SHJ17318.1_Bacteroides_stercorirosoris
WP_005879887.1_Oxalobacter_formigenes
WP_102737257.1_Akkermansia_muciniphila
WP_091534289.1_Alistipes_sp.
WP_027291455.1_Rikenella_microfusus
CCY64570.1_Prevotella_sp.
WP_032136094.1_Alistipes
ALJ40309.1_Bacteroides_thetaiotaomicron
DAB08223.1_Gastranaerophilales_bacterium
WP_047100952.1_Brachyspira_hyodysenteriae
WP_094324440.1_Campylobacter_avium
GM_11305_Giardia_muris
GM_20856_Giardia_muris
GM_24446_Giardia_muris
GM_21992_Giardia_muris
GM_21986_Giardia_muris
ESU41346.1_Giardia_intestinalis
XP_001306442.1_Trichomonas_vaginalis
XP_001321667.1_Trichomonas_vaginalis
WP_066104170.1_Gallibacterium_salpingitidis
WP_077473486.1_Rodentibacter
OHT15771.1_Tritrichomonas_foetus
OHT07803.1_Tritrichomonas_foetus
ODU03256.1_Pseudonocardia_sp.
WP_013676410.1_Pseudonocardia_dioxanivorans
WP_046732086.1_Streptomyces_humi
WP_097323795.1_Actinoplanes_atraurantiacus
WP_043441340.1_Arthrobacter_sp.
BAU96409.1_Corynebacterium_glutamicum
WP_068430993.1_Piscicoccus_intestinalis
WP_083163319.1_Mycobacterium_aquaticum
WP_081714872.1_Helicobacter_fennelliae
EFP61298.1_Erysipelotrichaceae_bacterium
WP_081341564.1_Agrobacterium_vitis
ACB91984.1_Xylella_fastidiosa
WP_058671364.1_Enterobacter_hormaechei
XP_003293280.1_Dictyostelium_purpureum
WP_066288176.1_Arthrobacter_sp.
KFF47432.1_Pseudomonas_sp.
WP_003807566.1_Bordetella_bronchiseptica
WP_076327646.1_Paenibacillus_amylolyticus
SFN22858.1_Psychrobacillus_psychrodurans
WP_106028886.1_Bacillus_pumilus
WP_103372469.1_Staphylococcus_nepalensis
CDA62234.1_Clostridium_sp.
ORX88120.1_Anaeromyces_robustus
OUM60671.1_Piromyces_sp.
WP_049974915.1_Blautia_schinkii
SCJ78981.1_Eubacterium_sp.
ORY22739.1_Neocallimastix_californiae
WP_087218163.1_Blautia_sp.
PGH24004.1_Fusobacterium_nucleatum
WP_102834397.1_Clostridioides_difficile
OBZ33202.1_Megasphaera_sp.
WP_094636049.1_Bifidobacterium_eulemuris
CDA18180.1_Acetobacter_sp.
ERJ30873.1_Campylobacter_concisus
WP_082987518.1_Gallibacterium_genomosp.
KXB49252.1_Corynebacterium_kroppenstedtii
WP_009142612.1_Succinatimonas_hippei
WP_082942594.1_Helicobacter
CDD81106.1_Dialister_sp.
WP_006915872.1_Lactobacillus_coleohominis
WP_075888181.1_Lactobacillus_crustorum
WP_010788710.1_Bacillus_atrophaeus
OJG89486.1_Enterococcus_raffinosus
WP_009302400.1_Desulfovibrio_sp.
WP_049687050.1_Dakarella_massiliensis
SNX32375.1_Lactobacillus_fermentum
WP_018371447.1_Streptococcus_massiliensis
WP_002607579.1_Clostridium
WP_071861943.1_Enterococcus_devriesei
CEI82750.1_Oceanobacillus_oncorhynchi
WP_006555587.1_Veillonella_seminalis
0.4

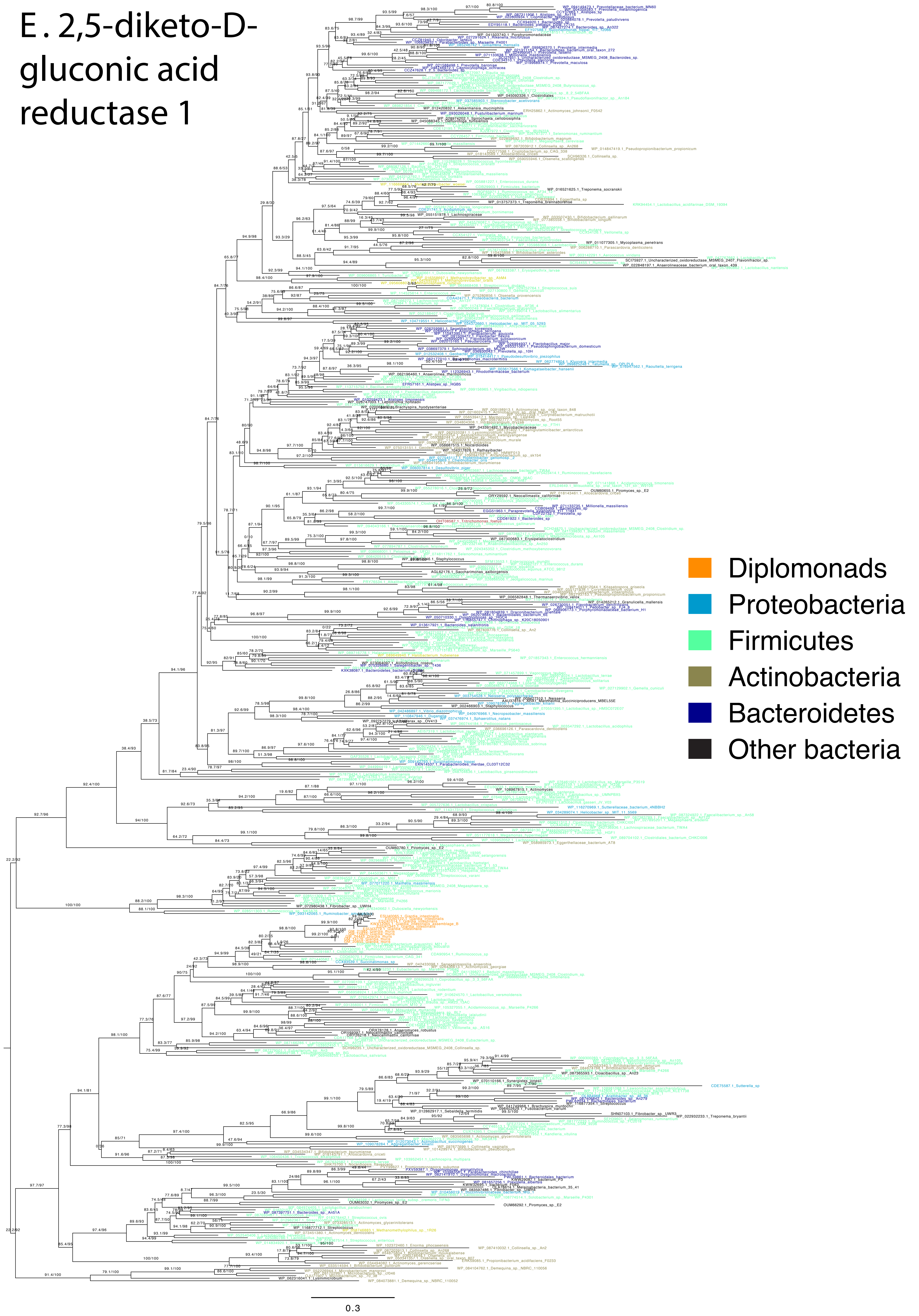
E. 2,5-diketo-D-gluconic acid reductase 1
Diplomonads
Proteobacteria
Firmicutes
Actinobacteria
Bacteroidetes
Other bacteria
0.3

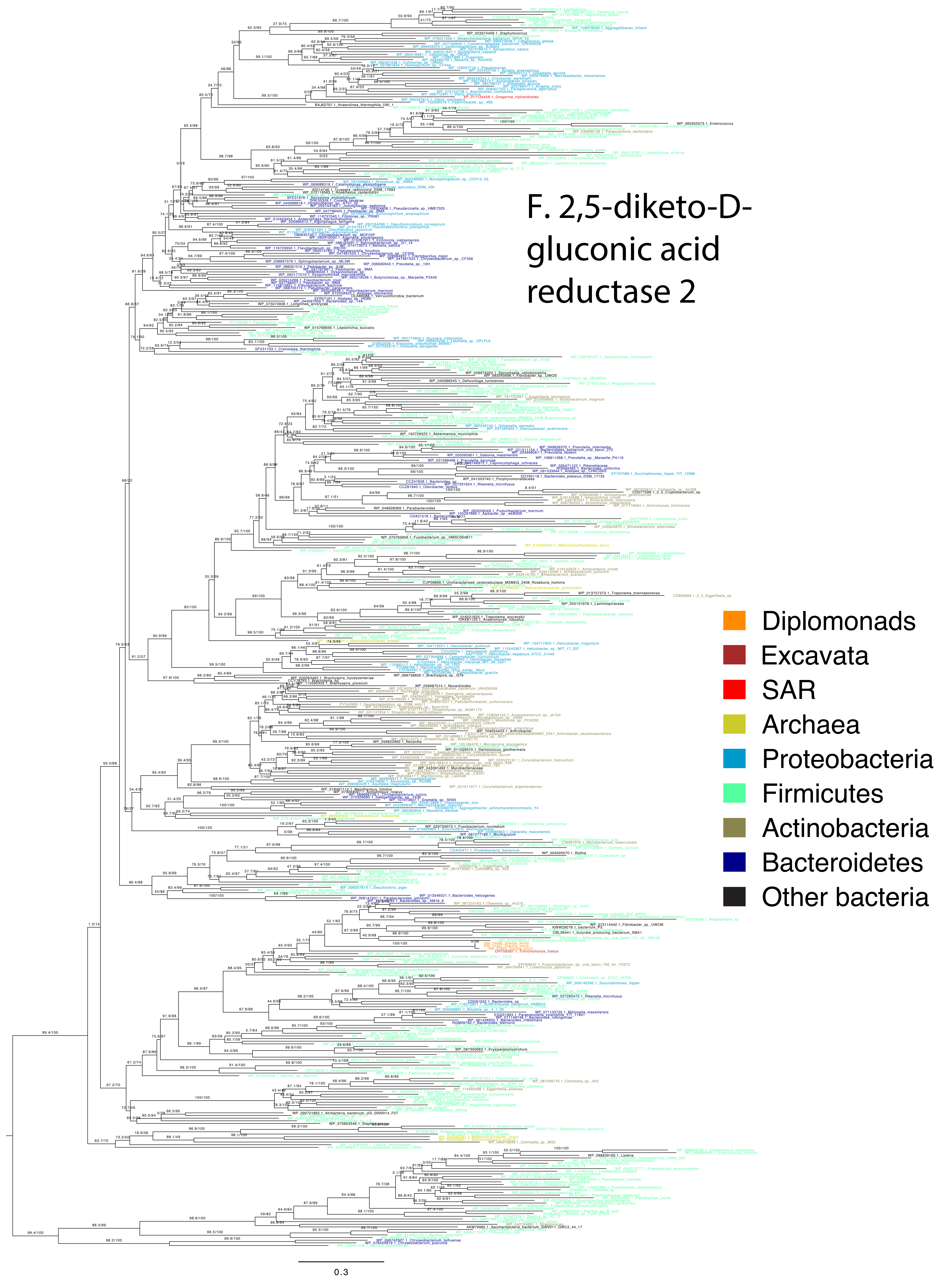
F. 2,5-diketo-D-gluconic acid reductase 2
Diplomonads
Excavata
SAR
Archaea
Proteobacteria
Firmicutes
Actinobacteria
Bacteroidetes
Other bacteria
0.3

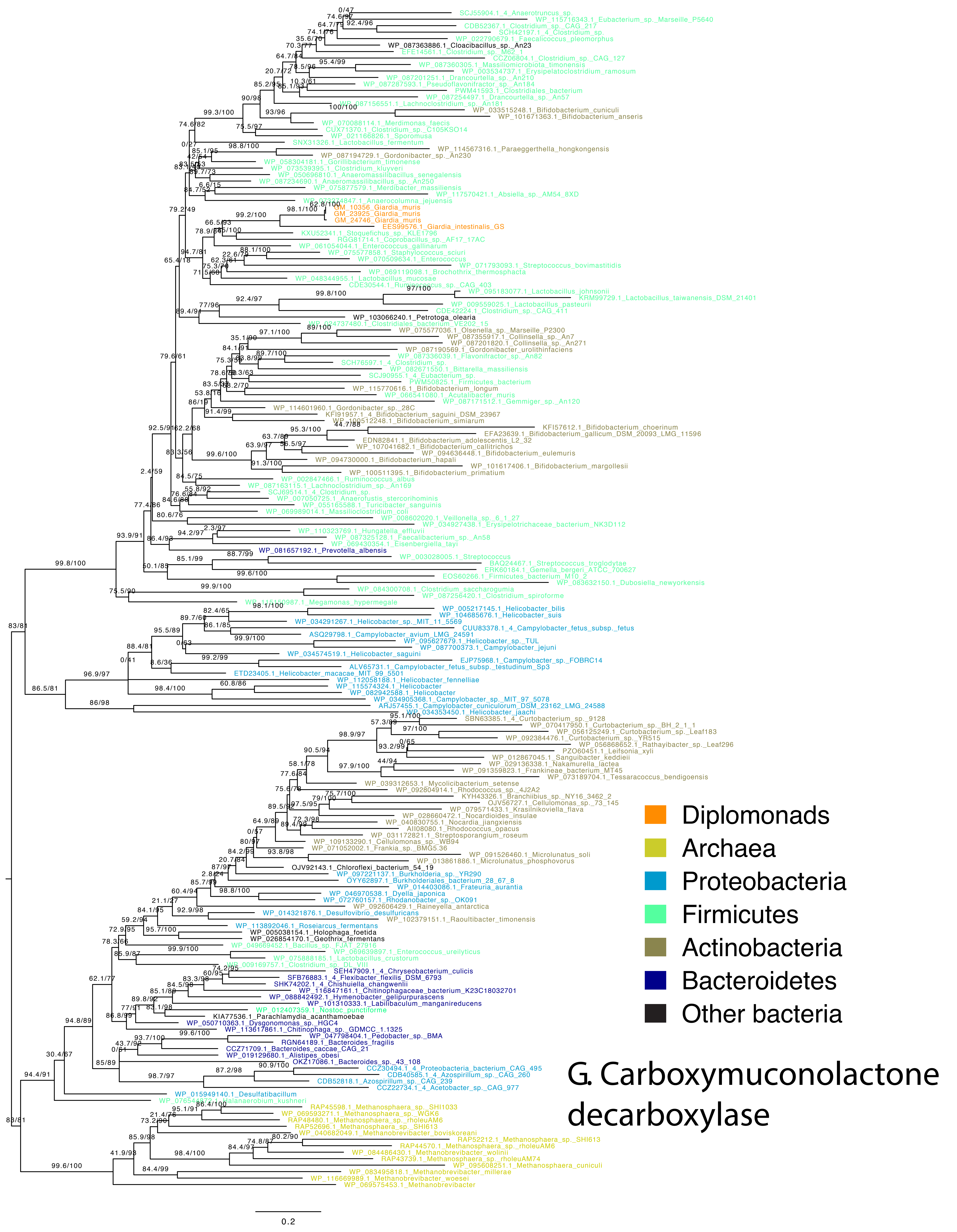

Diplomonads
Archaea
Proteobacteria
Firmicutes
Actinobacteria
Bacteroidetes
Other bacteria
G. Carboxymuconolactone decarboxylase
0.2

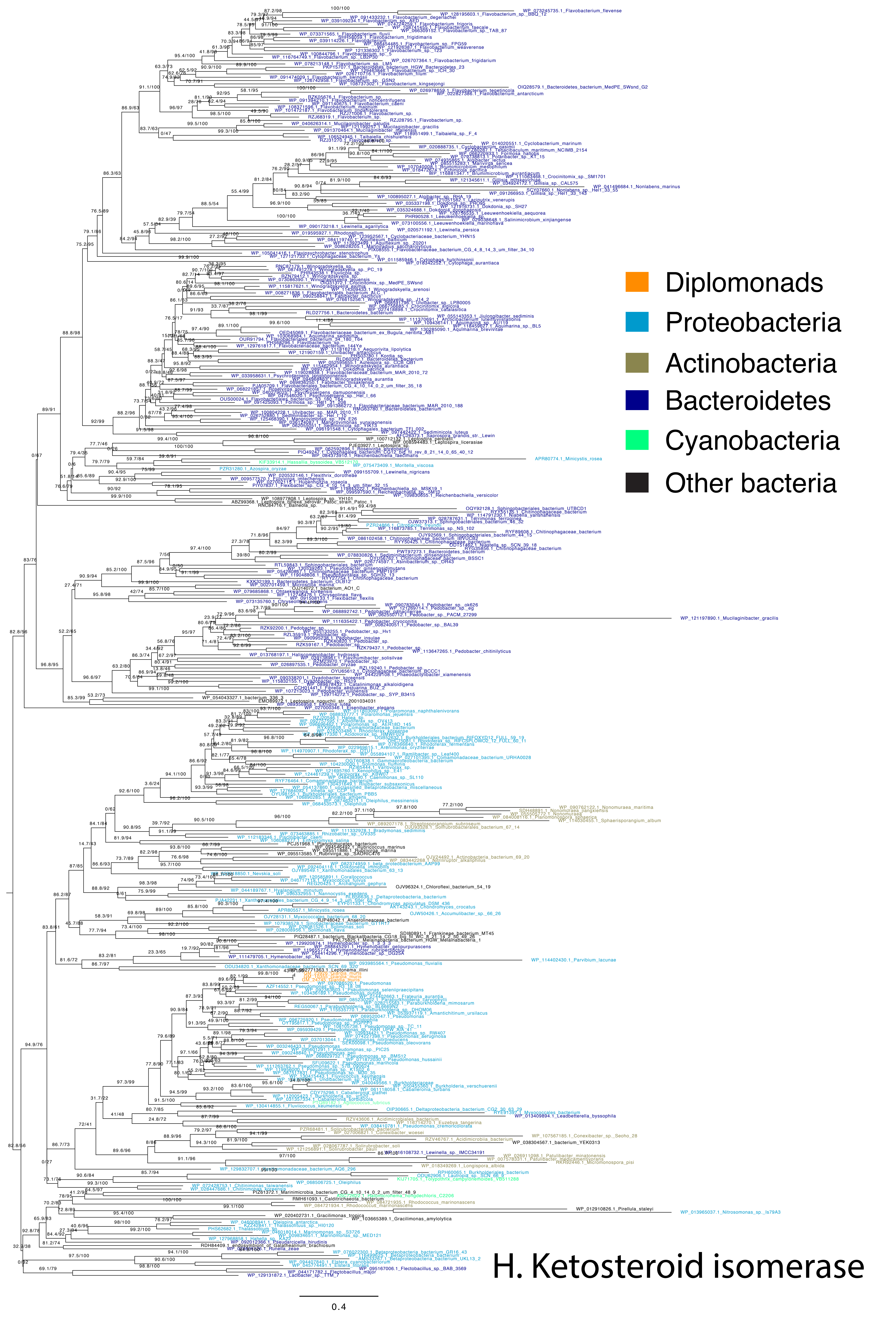


H. Ketosteroid isomerase

### I. Tryptophanase

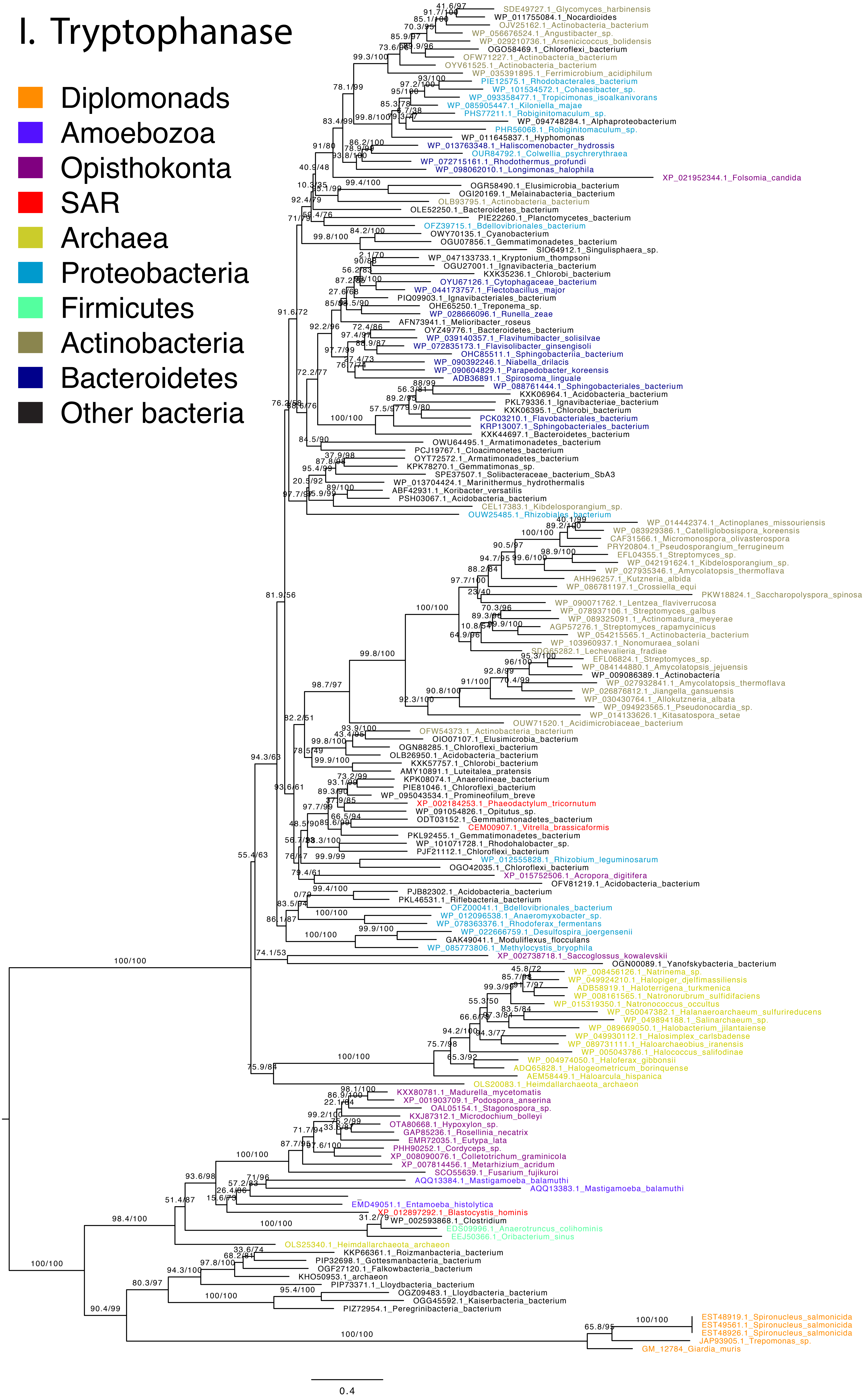

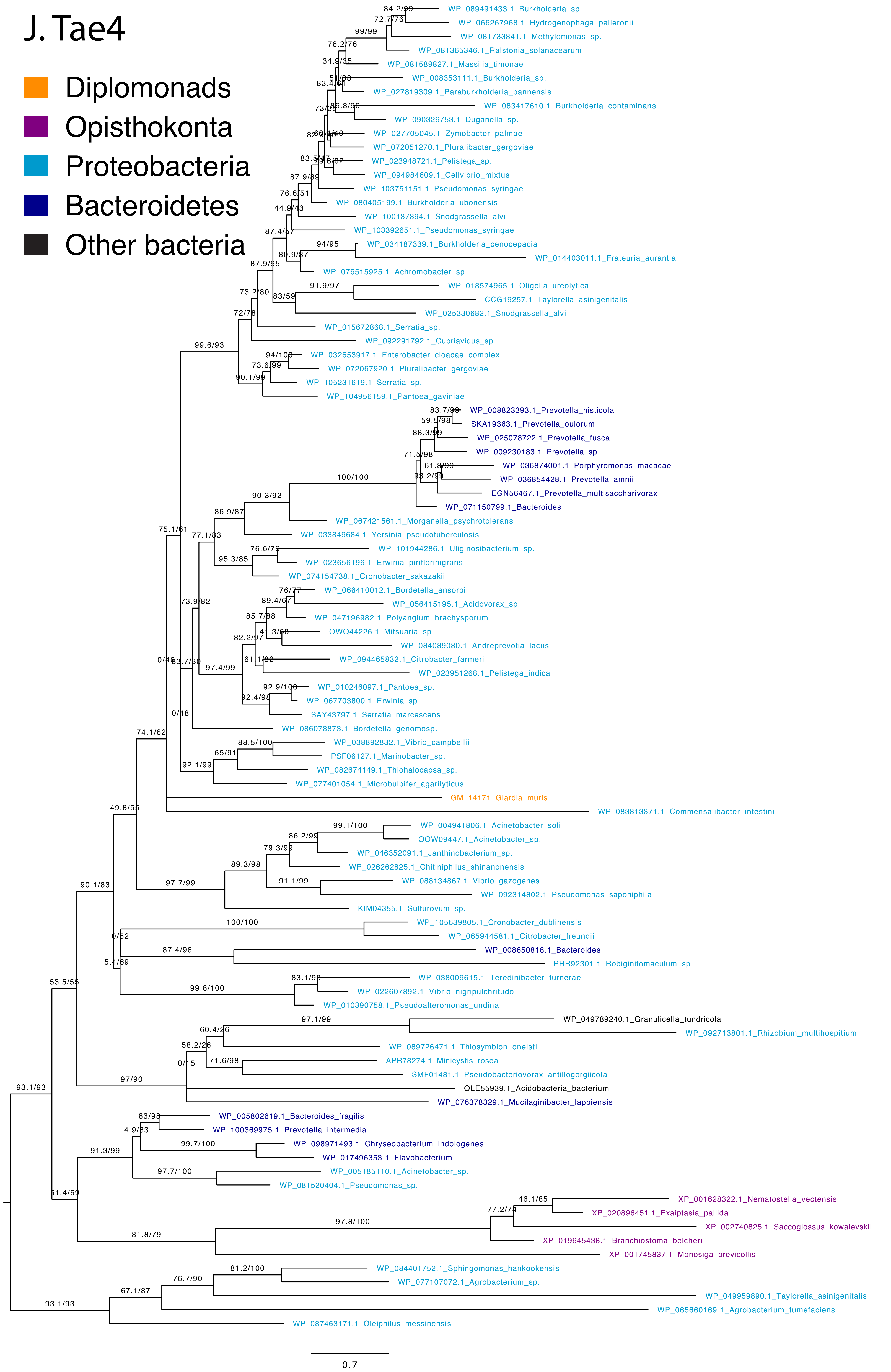

J. Tae4
Diplomonads
Opisthokonta
Proteobacteria
Bacteroidetes
Other bacteria
WP_089491433.1_Burkholderia_sp.
WP_066267968.1_Hydrogenophaga_palleronii
WP_081733841.1_Methylomonas_sp.
WP_081365346.1_Ralstonia_solanacearum
WP_081589827.1_Massilia_timonae
WP_008353111.1_Burkholderia_sp.
WP_027819309.1_Paraburkholderia_bannensis
WP_083417610.1_Burkholderia_contaminans
WP_090326753.1_Duganella_sp.
WP_027705045.1_Zymobacter_palmae
WP_072051270.1_Pluralibacter_gergoviae
WP_023948721.1_Pelistega_sp.
WP_094984609.1_Cellvibrio_mixtus
WP_103751151.1_Pseudomonas_syringae
WP_080405199.1_Burkholderia_ubonensis
WP_100137394.1_Snodgrassella_alvi
WP_103392651.1_Pseudomonas_syringae
WP_034187339.1_Burkholderia_cenocepacia
WP_014403011.1_Frateuria_aurantia
WP_076515925.1_Achromobacter_sp.
WP_018574965.1_Oligella_ureolytica
CCG19257.1_Taylorella_asinigenitalis
WP_025330682.1_Snodgrassella_alvi
WP_015672868.1_Serratia_sp.
WP_092291792.1_Cupriavidus_sp.
WP_032653917.1_Enterobacter_cloacae_complex
WP_072067920.1_Pluralibacter_gergoviae
WP_105231619.1_Serratia_sp.
WP_104956159.1_Pantoea_gaviniae
WP_008823393.1_Prevotella_histicola
SKA19363.1_Prevotella_oulorum
WP_025078722.1_Prevotella_fusca
WP_009230183.1_Prevotella_sp.
WP_036874001.1_Porphyromonas_macacae
WP_036854428.1_Prevotella_amnii
EGN56467.1_Prevotella_multisaccharivorax
WP_071150799.1_Bacteroides
WP_067421561.1_Morganella_psychrotolerans
WP_033849684.1_Yersinia_pseudotuberculosis
WP_101944286.1_Uliginosibacterium_sp.
WP_023656196.1_Erwinia_piriflorinigrans
WP_074154738.1_Cronobacter_sakazakii
WP_066410012.1_Bordetella_ansorpii
WP_056415195.1_Acidovorax_sp.
WP_047196982.1_Polyangium_brachysporum
OWQ44226.1_Mitsuaria_sp.
WP_084089080.1_Andreprevotia_lacus
WP_094465832.1_Citrobacter_farmeri
WP_023951268.1_Pelistega_indica
WP_010246097.1_Pantoea_sp.
WP_067703800.1_Erwinia_sp.
SAY43797.1_Serratia_marcescens
WP_086078873.1_Bordetella_genomosp.
WP_038892832.1_Vibrio_campbellii
PSF06127.1_Marinobacter_sp.
WP_082674149.1_Thiohalocapsa_sp.
WP_077401054.1_Microbulbifer_agarilyticus
GM_14171_Giardia_muris
WP_083813371.1_Commensalibacter_intestini
WP_004941806.1_Acinetobacter_soli
OOW09447.1_Acinetobacter_sp.
WP_046352091.1_Janthinobacterium_sp.
WP_026262825.1_Chitiniphilus_shinanonensis
WP_088134867.1_Vibrio_gazogenes
WP_092314802.1_Pseudomonas_saponiphila
KIM04355.1_Sulfurovum_sp.
WP_105639805.1_Cronobacter_dublinensis
WP_065944581.1_Citrobacter_freundii
WP_008650818.1_Bacteroides
PHR92301.1_Robiginitomaculum_sp.
WP_038009615.1_Teredinibacter_turnerae
WP_022607892.1_Vibrio_nigripulchritudo
WP_010390758.1_Pseudoalteromonas_undina
WP_049789240.1_Granulicella_tundricola
WP_092713801.1_Rhizobium_multihospitium
WP_089726471.1_Thiosymbion_oneisti
APR78274.1_Minicystis_rosea
SMF01481.1_Pseudobacteriovorax_antillogorgiicola
OLE55939.1_Acidobacteria_bacterium
WP_076378329.1_Mucilaginibacter_lappiensis
WP_005802619.1_Bacteroides_fragilis
WP_100369975.1_Prevotella_intermedia
WP_098971493.1_Chryseobacterium_indologenes
WP_017496353.1_Flavobacterium
WP_005185110.1_Acinetobacter_sp.
WP_081520404.1_Pseudomonas_sp.
XP_001628322.1_Nematostella_vectensis
XP_020896451.1_Exaiptasia_pallida
XP_002740825.1_Saccoglossus_kowalevskii
XP_019645438.1_Branchiostoma_belcheri
XP_001745837.1_Monosiga_brevicollis
WP_084401752.1_Sphingomonas_hankookensis
WP_077107072.1_Agrobacterium_sp.
WP_049959890.1_Taylorella_asinigenitalis
WP_065660169.1_Agrobacterium_tumefaciens
WP_087463171.1_Oleiphilus_messinensis
0.7

### K. Quorum-quenching N-acyl-homoserine lactonase

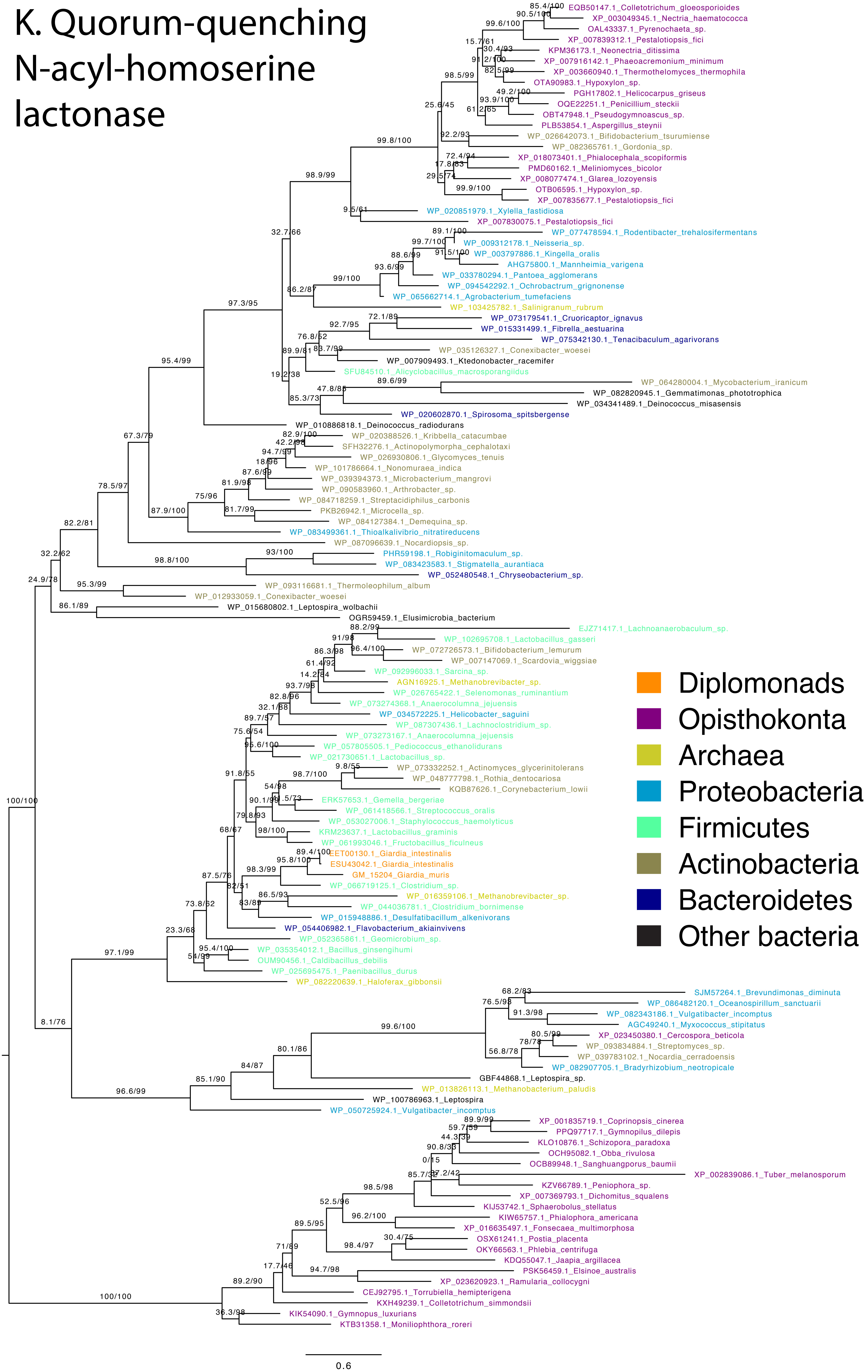
