## Supplementary material for "The compact genome of *Giardia muris* reveals important steps in the evolution of intestinal protozoan parasites": Methods S1

**DNA preparation from *G. muris* for long read sequencing**

*G. muris* excyzoites (~9×10^7^ cells) were centrifuged at 500 ×g for five minutes to remove the excystation media and the pellet was dissolved in Lysis Buffer (50 mM EDTA, 1% SDS and 10 mg/ml Proteinase K) by vortexing. The lysed cells were incubated at 56 °C with occasional vortexing for 2 hours followed by addition of 1 µl RNase A (100 mg/µl) and incubation at room temperature for 10-15 minutes. The genomic DNA was extracted by addition of 0.5 volumes each of phenol and CHISAM (chloroform: isoamyl alcohol 24:1). The sample was mixed carefully by vortexing and centrifuged at 13,000 rpm for 10 minutes, the water phase was collected and an equal volume of CHISAM was added to the sample and mixed by vortexing followed by a second centrifugation with same settings. The water phase was collected, and the DNA was precipitated with an equal volume of isopropanol. The precipitate was recovered by immediate centrifugation at 13,000 rpm, 4 °C, 30 min. The supernatant was removed, and the pellet was washed with 70% ice cold ethanol and then centrifuged at 13,000 rpm, 4 °C, 10 minutes. The supernatant was removed, air-dried briefly and resuspended in TE-buffer (10 mM Tris-HCl pH 8.0, 1 mM EDTA). The genomic DNA was further purified using Genomic-tip 20/G (Qiagen) according to the manufacturers instructions.

Finally, the genomic DNA was purified using phenol-chloroform extraction according to the “Extracting DNA Using Phenol-Chloroform” protocol (https://www.pacb.com/wp-content/uploads/2015/09/Experimental-Protocol-Extracting-DNA-using-Phenol-Chloroform.pdf). The DNA solution was mixed with an equal volume of the phenol/chloroform/isoamyl alcohol solution (25:24:1). The tube was mixed vigorously for 1 min and spun at high speed for 5 min. The aqueous phase was removed, and the tube was back extracted using EB buffer. The original and back-extracted material was combined and extracted as above using chloroform/isoamyl alcohol (24:1). The aqueous phase was precipitated using 2.5× volume 100% ethanol and NH_4_OAc (0.75 M final concentration) while adding glycogen (20 ug /extraction). The precipitate was recovered by high speed centrifugation and washed two times by double volumes of 80% ethanol. The pellet was air-dried and brought up in a small volume of EB buffer. The concentration and quality of the extracted DNA was determined by NanoDrop and agarose gel electrophoresis.

**RNA purification**

Total RNA from trophozoites were extracted using TriZol according to the method in Franzén et al.(1). The total RNA was DNase treated using the Turbo DNA-free kit (Applied Biosystems, AM1907) as described in Franzén et al.(1). RNA from cysts and excyzoites were extracted using TriZol as described above and the resulting RNA was DNase treated (Turbo DNase, 2 U/µl) in a reaction volume of 100 µl for 30 minutes at 37°C. The RNA was subsequently cleaned using the cleanup protocol from the RNeasy Mini Kit (Qiagen, 74104) and eluted in 30 µl of DEPC treated water. The quality and quantity were evaluated using Qubit and BioAnalyzer.

**Sequencing and assembly**

Total genomic DNA was sequenced using both Illumina MiSeq and PacBio RS II sequencers at the SNP&SEQ platform and Uppsala Genome Center respectively at the Science for Life Laboratory (Uppsala University). The trophozoite total RNA was divided in two parts and stranded transcriptome mRNA library was prepared using the appropriate TruSeq kit at GATC Biotech (Konstanze, Germany). The finished libraries were sequenced using the Illumina HiSeq 2000 system at GATC Biotech (Konstanz, Germany). RNA sequencing libraries from cysts and excyzoites were prepared using the TruSeq stranded mRNA sample preparation kit and sequenced by HiSeq 2500 at the SNP&SEQ platform.

Raw DNA and RNA sequence reads are archived at NCBI Sequence Read Archive (SRA) under accession numbers SRR8858297-SRR8858305.

**Biosamples and accession numbers**

| Biosample accession | Sample description | DNA/RNA | Sequencing instrument | Usage | SRA accession |
| --- | --- | --- | --- | --- | --- |
| SAMN11231833 | Cysts | DNA | PacBio RS II | Genome assembly | SRR8858302 - SRR8858305 |
|  |  | RNA | Illumina HiSeq 2500 | RNA-Seq comparison | SRR8858298 |
| SAMN11231832 | Trophozoites | DNA | Illumina MiSeq | Assembly base correction | SRR8858300 |
|  |  | RNA | Illumina HiSeq 2000 | RNA-Seq comparison | SRR8858301 |
| SAMN11231834 | Excysted trophozoites | RNA | Illumina HiSeq 2500 | RNA-Seq comparison | SRR8858299 |

**Genome assembly and contamination removal**

PacBio long reads alone were used for *de novo* genome assembly using the standard SMRT Analysis (v2.3.0) pipeline. Reads were assembled with HGAP followed by consensus sequence calling with Quiver. The resulting assembly contains 317 contigs. A BLASTN (evalue <= 0.1) of the contigs against the nt database revealed contamination, mostly from fungi, and the contaminated contigs were removed from the final assembly, resulting in 59 final contigs.

The Illumina MiSeq reads were mapped to the PacBio assembly using BWA v0.7.12-r1039 (2) and Nesoni v0.130 (http://bioinformatics.net.au/software.nesoni.shtml) was used to correct mostly the indels that we have observed to cause frame-shifts in certain genes. 8 deletion, 46 insertion and 16 SNPs were corrected by nesoni with setting --majority 0.75 based on the mapped bam file.

This Whole Genome Shotgun project has been deposited at DDBJ/EMBL/GenBank under the accession PRJNA524057. The version described in this paper is version VDLU00000000.1.

**Repeat detection**

RepeatMasker v3.3.0 (<http://www.repeatmasker.org/>) was run to detect genome repeats. Sequence comparison in RepeatMasker was performed by cross_match v1.080812. RM database v20110920 with RepBase Update 20110920 was used as RepeatMasker library.

**Heterozygosity estimation**

Samtools mpileup with B flag was used to generate pileup file from MiSeq reads mapped bam file. Sites with at least 20X coverage and 10% of alternative bases were considered as allelic heterozygous sites.

**Genome annotation**

Two gene prediction programs were used, Prodigal v2.60 and GlimmerHMM v3.0.1. Training set for GlimmerHMM consists of genes predicted from Prodigal which have hits against *G. intestinalis* with evalue <1e-50 and similar length to the hit gene. The union of the two sets of predicted genes was taken for functional annotation. Genes were manually annotated using information from BLASTP results against NR database as well as HMMER (v3.0) search results of domain information against Pfam (v27.0).

**tRNA and rRNA**

18S, 28S and 5.8S ribosomal RNAs were annotated after BLASTN similarity to the deposited *G. muris* ribosomal RNAs in NCBI (X65063), and the start and stop positions were adjusted to be inline with the record. 5S rRNAs were predicted by searching against Rfam 11.0 using infernal v1.0.2. There are 78 pieces of 28S rRNAs, 73 18S rRNAs, 68 5.8S rRNAs and 10 5S rRNA annotated.

tRNAs were predicted by tRNAScan-SE v1.23 with the most sensitive co-variance model.

BLASTN with -task blastn (which is more sensitive for distant matches) of the 40 *G. intestinalis* ncRNAs against *G. muris* genome revealed 10 ncRNAs with evalue <0.1 and not overlapping genes. Alignment of the 10 ncRNAs using MUSCLE v3.8.31 showed a clear motif on the 3’ end, CCTTYNHTNAA, which is similar to what was found on *G. intestinalis* (CCTTYNHTHAA). This motif was used to search for other potential ncRNAs in *G. muris* using scan_for_matches (http://blog.theseed.org/servers/2010/07/scan-for-matches.html). Additionally, results were obtained from BLASTN search against miRBase, similarity search against Rfam and prediction of small RNAs based on small RNA-Seq data using Shortstack v1.2.4. All the search and prediction results were combined following the rule that ncRNAs do not overlap (or overlap little) with the annotated genes and with themselves. Manual efforts were also applied. In the end, they are 219 ncRNAs annotated, with 16 from similarity search against *G. intestinalis* ncRNAs, 8 from Rfam searches, 17 from Shortstack prediction, and 178 from motif search.

**Protein kinases**

Protein kinases were assigned into group, family and subfamily. Alignment files for different group, family and subfamily were downloaded from the kinase database (<http://kinase.com>). HMMER 3.0 (<http://hmmer.janelia.org>) was used to build HMM profiles from the alignment and the protein sequences were searched against the HMM profiles. The search results were then combined with Pfam domain search results where genes contain significant Pfam Pkinase domain (PF00069) (score >25). *G. intestinalis* protein kinase annotations were taken from reference.

**Ankyrin repeat proteins**

Ankyrin repeat proteins (ARPs) were assigned to genes that contain ankyrin repeat domains (score >=20) but not Pkinase domain. According to if an ARP contains zink finger domains or other domains, they were further divided into 3 subgroups.

**Cysteine-rich proteins**

Alignments of the proteins with more than 10% cysteines using MAFFT v7.215 revealed that a subset of them contains a conserved pentapeptide motif (GR)CR(GRKE)K at the C-terminus. Cysteine-rich proteins were divided into three subgroups depending on the presence and absence of the transmembrane domains and the pentapeptide motif.

**Synteny**

MUMmer v3.23 was used to align the draft genomes of *G. muris* and *G. intestinalis*. Promer was used in order to align the genome at protein level. Mummerplot was used to generate the alignment dot plot. Show-coords was used to view a summary of all the alignments produced by promer. Alignment coordinates were used to calculate the percentage of similarity between the two genomes as well as to draw the similarity in Circos plot.

OrthoMCL v2.0.2 was used to cluster genes using match cutoff of 40% and e-value cutoff of 1e-10. The clustering was done within *G. muris*, between *G. muris* and *G. intestinalis*, and among *G. muris*, *G. intestinalis* and *S. salmonicida*.

**RNA-Seq expression analysis**

BAM files were generated from mapping the RNA-Seq reads to the reference genome using BWA v0.7.12-r1039. Cufflinks v104700 was used to calculate the FPKM values from the BAM files. FPKM by definition: # read pairs in genes / # total reads / 1,000,000 / size of the gene.

**Phylogenetic trees for NEKs and ARPs**

All the NEKs from *G. muris* and *G. intestinalis* were aligned with MAFFT v7.407 (auto setting), and FastTree v2.1.10 No SSE3 was used to construct approximately-maximum-likelihood tree using default WAG+CAT model. ARPs tree was constructed the same way as NEKs tree.

**Phylogenetic analysis of lateral transfers**

One sequence from *G. muris* was used as a query for each enzyme to retrieve at least 5000 hits with e-value <0.001, using BLASTP against the nr database and the organism-specific proteomes. The resulting datasets were filtered automatically by finding all sequences with greater than 90% sequence identity to another sequence using the software skipredundant (EMBOSS v6.5.7.0 (3)) with default settings. Datasets were aligned using MAFFT v6.603b (4), and regions of ambiguous alignment were automatically trimmed using BMGE v1.12 (blosum30 and block size of 2) (5). Sequences with a length less than 50% of the multiple alignment were removed, and the datasets were aligned and trimmed again as described above. An initial phylogenetic analyses were performed using FastTree v2.1.8 SSE3, OpenMP (6). Sequences with a phylogenetic distance < 0.3 were removed in an iterative process to create the final datasets. This final datasets were aligned in forward and reverse orientation using MAFFT and PROBCONS v1.12 (7). The four resulting alignments were combined with T-COFFEE v11.00.8cbe486 (8) into a maximal consensus alignment, which was trimmed using BMGE as in previous steps. Maximum Likelihood (ML) trees were computed using IQtree v1.6.5 (9) under LG4X substitution model (10). Branch supports were assessed using ultrafast bootstrap approximation (UFboot) with 1,000 ultrafast bootstraps (11) and 1,000 replicates for SH-like approximate likelihood ratio test (SH-aLRT) (12).

**Promoter motif search**

MEME suite v4.10.2 was used for promoter motif analysis. A maximum of 120 bp upstream of annotated genes were used in search of potential promoter. The longest possible sequence was used if the intergenic region is shorter than 120 bp, and the sequence was discarded if the intergenic region is shorter than 8 bp. 4139 sequences in total were used in motif search, and MEME was set to search for 10 most likely motifs with sizes from 6 to 30 bp.

**VSP and ψVSP phylogenetic analysis**

Nucleotide sequences of *G. muris* VSPs and ψVSP genes were aligned using MAFFT in MAFFT v7.416(4), and regions of ambiguity were trimmed using BMGE v1.12 (DNAPAM100:2, (5)). Maximum Likelihood (ML) trees were computed using IQtree v.1.6.5 (9) under the best model (TIM2+F+R6) with 1,000 ultrafast bootstraps (Hoang et al., 2018) and 1,000 bootstrap replicates for SH-like approximate likelihood ratio test (SH-aLRT) (12).
